# ECMExplainer: a functional matrisome framework for decoding cellular stress states and aging trajectories

**DOI:** 10.64898/2026.11.11.766024

**Authors:** Eleanor Sheekey, Roberto A. Avelar, Hironobu Fujiwara

## Abstract

The extracellular matrix (ECM) is increasingly recognized as both a hallmark of aging and a durable record of the cellular stress states that shape age-related tissue dysfunction. Despite its apparent role in aging and disease, translating matrisome gene expression into interpretable biological meaning remains an unresolved challenge. Here we introduce ECMExplainer, a framework that uses 180 matrisome gene-pair ratios, partitioned into 76 Assembly/Destruction and 104 Signaling/Signaling pairs to position fibroblast transcriptomes within a functional matrisome phenotype space defined by six reference fibroblast states spanning physiological and stress-associated conditions: Proliferating, Quiescence, Senescence, Fibrosis, Inflammation, and Hypoxia. Because each ratio records only which of the two genes is more highly expressed, it is largely insensitive to sequencing depth and batch offsets, allowing ECMExplainer to outperform magnitude-based classifiers on unseen data. Rather than assigning only a single phenotype label, ECMExplainer measures a sample’s coordinate distance to each reference state, enabling quantitative mapping of matrisome state trajectories while giving biological meaning even to conditions that fall between or outside the training phenotypes. Across 20 independent fibroblast datasets, ECMExplainer uncovered conserved ECM signatures underlying diverse fibroblast states and showed that they capture multiple aging-related processes. Specifically, it identified FST/ITGA1 as a rapamycin-sensitive senescence-associated ratio, revealed a fibrosis-biased ECM trajectory in Hutchinson-Gilford progeria syndrome distinct from chronological aging, and distinguished successful from failed OSKM reprogramming using an ECM Aging Score without retraining. Together, these findings establish the matrisome as a compact and biologically interpretable carrier of cellular state information, and ECMExplainer as a quantitative foundation for interpreting aging, fibrosis and rejuvenation through the lens of the ECM.

**Graphical abstract:** 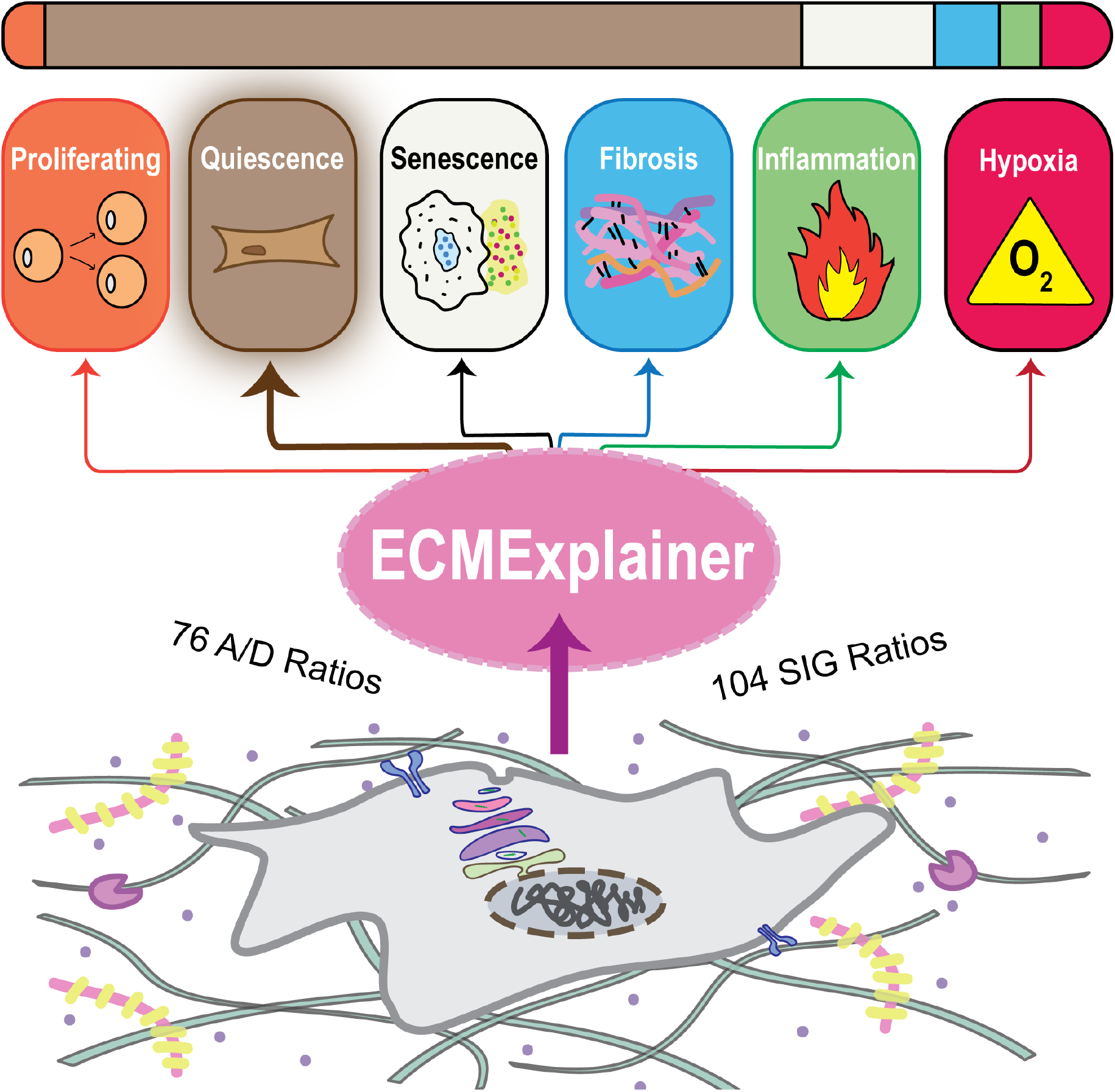

## 2 Introduction

Because cells continuously construct, remodel, and respond to their extracellular matrix (ECM), the ECM serves not only as a structural scaffold but also as a relatively stable repository of information about cellular and tissue state. Deciphering this information could provide a unique window into the reciprocal communication between cells and their extracellular environment. According to the MatrisomeDB, the human ECM comprises over 1,000 genes encoding structural matrisome components, secreted factors involved in ECM crosslinking, degradation, and remodeling, as well as proteins that mediate tissue-level and systemic communication ^1–3^. The expression of these genes differs by tissue and cell type, together contributing to homeostatic organ function. With the advent of new experimental tools and novel mouse models ^4^, our understanding of individual ECM protein functions is expanding rapidly. To date, however, much of our knowledge of the ECM is organized at the level of individual molecules, their binding partners, and their tissue distributions. This molecular catalogue is a necessary foundation, but it does not by itself explain how diverse ECM components collectively encode tissue state or regulate cellular behavior. Ultimately, understanding the ECM requires a functional framework that interprets ECM composition as an integrated representation of cell–ECM interactions, rather than as a collection of individual molecules.

This functional gap is particularly relevant to aging. Matrix alterations, including changes in stiffness, composition, and turnover, have recently been proposed as a hallmark of the process ^5,6^, and a plasma proteomic clock constructed from only 14 circulating ECM proteins can accurately predict chronological age and distinguish healthy from diseased states ^7^. A defining feature of aged tissue is the progressive accumulation of senescent cells, which persist long after the stressors that triggered them, secreting a senescence-associated secretory phenotype (SASP) that reshapes the surrounding matrix ^8^. In addition, tissue fibrosis increases with age, resulting in elevated collagen deposition, enhanced crosslinking, and impaired matrix turnover ^9^. Fibroblasts within an aged tissue can span a range of states from proliferative through to senescent or fibrotic, each imposing a characteristically different ECM program. The aged matrisome therefore reflects a composite of coexisting cellular phenotypes rather than a single "old" signature. Critically, the aging ECM is not merely a downstream product of these cellular states but can act upstream of them. Age-associated changes in stiffness, composition, and ligand presentation modulate integrin engagement and mechanosensing, biasing damaged cells toward senescence and supporting the persistence of established senescent cells which in turn remodel the matrix further. This forms a self-reinforcing circuit that amplifies tissue aging ^10^. Because ECM remodeling is thus both a consequence and a driver of aging, the matrisome gene expression captures a distinct layer of biological information that is not encoded by cell-intrinsic transcriptional programs alone. Rather than describing only the state of individual cells, it reflects the extracellular environment that cells collectively build, remodel, and interpret. Realizing that potential, however, is not simply a matter of cataloguing which genes change. Many studies have documented that ECM gene expression shifts with age, yet demonstrating that the matrisome changes is distinct from interpreting what those changes mean for cellular states. Indeed, the latter remains unresolved because the field still lacks a functional framework to translate an age-altered ECM expression profile into the phenotypic programs it represents.

Despite these clear phenotypic links, detection of states such as fibrosis in practice often relies on measuring highly abundant structural ECM components such as through Masson’s trichrome staining or the quantification of Collagen Type I (COL1A1) expression ^11^. As these structural proteins are already highly expressed in healthy tissues, relying on them as primary markers yields a poor signal-to-noise ratio for early or subtle phenotypic shifts. Furthermore, absolute expression of a single component cannot capture the dynamic state of the matrisome. Several frameworks have begun to decode context-specific changes in ECM gene-expression programs. A “Collagen Code” was proposed to capture alterations in the tumor microenvironment ^12^. Similarly, Collagen Disorder Architecture (CoDA), which quantifies collagen fragmentation, bundling, anisotropy, density, and rigidity, has been used to stratify tumors by stage, molecular subtype, and mutational status ^13^. Beyond cancer, an aging-related counterpart emerged through the definition of “matreotypes”, matrisome gene-expression patterns associated with age, derived from age-stratified human transcriptomes ^14^. In parallel, broader matrisome-focused tools have also been developed: Matrisome AnalyzeR provides standardized matrisome annotation across organisms ^15^; MatriCom infers cell–ECM and ECM–ECM interactions from scRNA-seq data ^16^; and most recently, MatriSpace ^17^ enables visualization and quantification of matrisome gene-set expression across Visium datasets spanning healthy and diseased tissues. Collectively, these approaches have advanced the descriptive, organizational, and spatial characterization of ECM biology. However, a complementary phenotypic framework remains lacking, one capable of taking an ECM gene-expression profile from any platform and positioning the sample within a functional ECM state space encompassing multiple homeostatic and stress phenotypes. Such a framework could additionally quantify how interventions shift samples along each phenotypic axis.

To build such a framework, we reasoned that, despite its molecular complexity, the primary functions of the matrisome can be abstracted into three fundamental processes: Assembly genes encode the structural proteins and factors that build and stabilize the ECM. Destruction genes encode the enzymes that degrade it. Lastly, Signaling genes include the receptors and secreted factors that sense, transmit, and coordinate extracellular cues, thereby regulating both ECM homeostasis and cellular or tissue states. Assigning each matrisome gene to one of these three functional modules, Assembly (A), Destruction (D), and Signaling (SIG), reframes a large, heterogeneous gene set in terms of the biological process it contributes to, and captures the idea that ECM state is determined by the dynamic interplay between matrix assembly, matrix destruction, and the signaling processes that coordinate these processes while mediating communication between cells and their extracellular environment. Building on this logic, we rationalized that gene ratios would quantify matrix states more effectively than absolute expression levels because they represent the relative balance among complementary matrisome functions while remaining intrinsically robust to platform, library-size, and batch effects (which cancel within the ratio) and provide a more sensitive readout of phenotypic state.

Here, we computationally evaluated the discriminative power of all possible A/D and SIG/SIG gene-pair ratios across six well-defined phenotypic states: Proliferation, Quiescence, Senescence, Fibrosis, Inflammation, and Hypoxia. This mathematically unbiased search identified highly discriminative, and often unexpected, ECM gene targets, and yielded ECMExplainer, a lineage-agnostic and intrinsically batch-robust framework that translates complex ECM gene-expression profiles into interpretable functional states. ECMExplainer is best understood not as a conventional single-label classifier but as a coordinate system for matrisome state. Rather than forcing every sample into one of the six categories, the ratios that separate the reference phenotypes together define a continuous phenotype space in which any expression profile can be positioned and its distance to each reference state measured. This gives biological meaning even to samples that fall between or outside the training phenotypes. More broadly, ECMExplainer provides a general strategy for decoding the functional information contained in matrisome expression profiles across diverse biological contexts. We demonstrate this utility across aging datasets, partial reprogramming, and the quantitative mapping of rapamycin-mediated ECM rescue, revealing continuous transitions between extracellular functional states.

## 3 Results

### 3.1 Training the ECMExplainer

Fibroblasts adopt distinct phenotypic states each thought to be accompanied by characteristic remodeling of the surrounding extracellular matrix (ECM). To ask whether these states could be resolved directly from ECM-focused gene expression, we first required a gene panel that represents the full life cycle of the matrix rather than any single compartment. We therefore set out to define a panel that jointly captures the three core processes by which fibroblasts establish and maintain their matrix: (i) Assembly, the synthesis, post-translational processing, and supramolecular organization of matrix components; (ii) Destruction, the proteolytic and glycolytic turnover that remodels the matrix; and (iii) Signaling, the receptors and secreted factors that modulate communication between cells and the matrix, enabling cells to sense, interpret, and respond to extracellular cues. To keep the panel ECM-specific and avoid contamination by broad transcriptional or cell-state programs, we built it exclusively from curated, ECM-focused gene sets and admitted a gene only if it was a member of at least one such set. We refer to this curated resource as the ECMExplainer gene list throughout.

We assembled the ECMExplainer gene panel from the Human Matrisome Portal masterlist (Naba et al., 2012) together with nine curated ECM-focused pathway gene sets from MSigDB (v2026.1.Hs), representing annotations from complementary knowledge bases including Reactome, Gene Ontology, and KEGG (Liberzon et al. 2015; see Methods). Together these resources encompass structural matrisome components, ECM assembly and degradation pathways, and cell–ECM signaling. After harmonization to current HGNC gene symbols and limited manual curation to remove genes lacking clear ECM relevance in fibroblasts (Methods), the final ECMExplainer panel comprised 1,139 genes assigned to three functional modules: Assembly (n = 412), Destruction (n = 119), and Signaling (n = 608) (Supplementary Table 1).

To identify ECM genes with the greatest ability to distinguish cellular phenotypes, we curated publicly available RNA-sequencing datasets from human fibroblasts. Ten studies spanning 96 samples across six distinct cellular phenotypes (Proliferating, Senescence, Quiescence, Fibrosis, Inflammation and Hypoxia, Supplementary Table 2) were selected to cover key fibroblast contexts relevant to aging and disease ^19,20^. Cell lines included fetal human skin and lung fibroblasts (BJ, HFF, IMR90, WI-38), primary human lung fibroblasts, and induced pluripotent stem cell-derived cardiac fibroblasts (iPSC-CFs) (Supplementary Table 2), covering distinct anatomical origins and developmental stages rather than a single matched lineage. As ECM composition and gene expression are strongly tissue-specific ^2^, we intentionally incorporated multiple tissue origins during training to reduce the risk that the model learns tissue- or age-specific expression patterns rather than phenotype-generalizable ECM signatures. For each training dataset, PCA was performed independently to confirm phenotypic separation within each study prior to integration (Supplementary Figure 1). The raw expression matrices were then merged across studies. To verify that all samples represented a common fibroblast lineage despite their diverse phenotypic states, each of the 96 samples was scored against a canonical fibroblast marker panel^21,22^ and against epithelial, endothelial, immune, smooth-muscle and pluripotent-stem-cell lineage panels drawn from the Tabula Sapiens human cell atlas ^23^. Smooth-muscle markers were further restricted to the contractile-phenotype set defined by Owens et al. 2004, and pluripotency was assessed using the canonical reprogramming factors ^25^. Fibroblast identity was the dominant lineage score in all 96 samples, whereas non-fibroblast lineage scores remained uniformly low (Supplementary Figure 2A). Smooth-muscle markers were elevated specifically in the Fibrosis samples of GSE102674, consistent with myofibroblast activation during TGFβ-driven fibrotic induction ^26^ rather than non-fibroblast contamination. Global PCA of the uncorrected matrix showed samples clustering primarily by study rather than phenotype for both the top 2,000 variable genes and the curated ECM gene list, indicating a strong batch effect requiring correction prior to phenotype-level analysis (Supplementary Figure 2B).

Expression matrices were therefore batch-corrected using limma prior to downstream analysis. Genome-wide PCA of the corrected data separated the six phenotypes (PC1 25.2%, PC2 14.7%; Figure 1A), recovering the biological structure previously obscured by study-specific batch effects in Supplementary Figure 2B. Of 1,139 curated ECMExplainer genes, 895 passed quality filtering and were retained for all subsequent ECM-space analyses (Figure 1B). PCA restricted to these 895 ECM genes further sharpened phenotypic separation, most notably resolving Senescence from the remaining phenotypes (PC1 21.1%, PC2 13.5%; Figure 1B), indicating that fibroblast cell states are specifically encoded within the ECM transcriptome.

**Figure 1:**
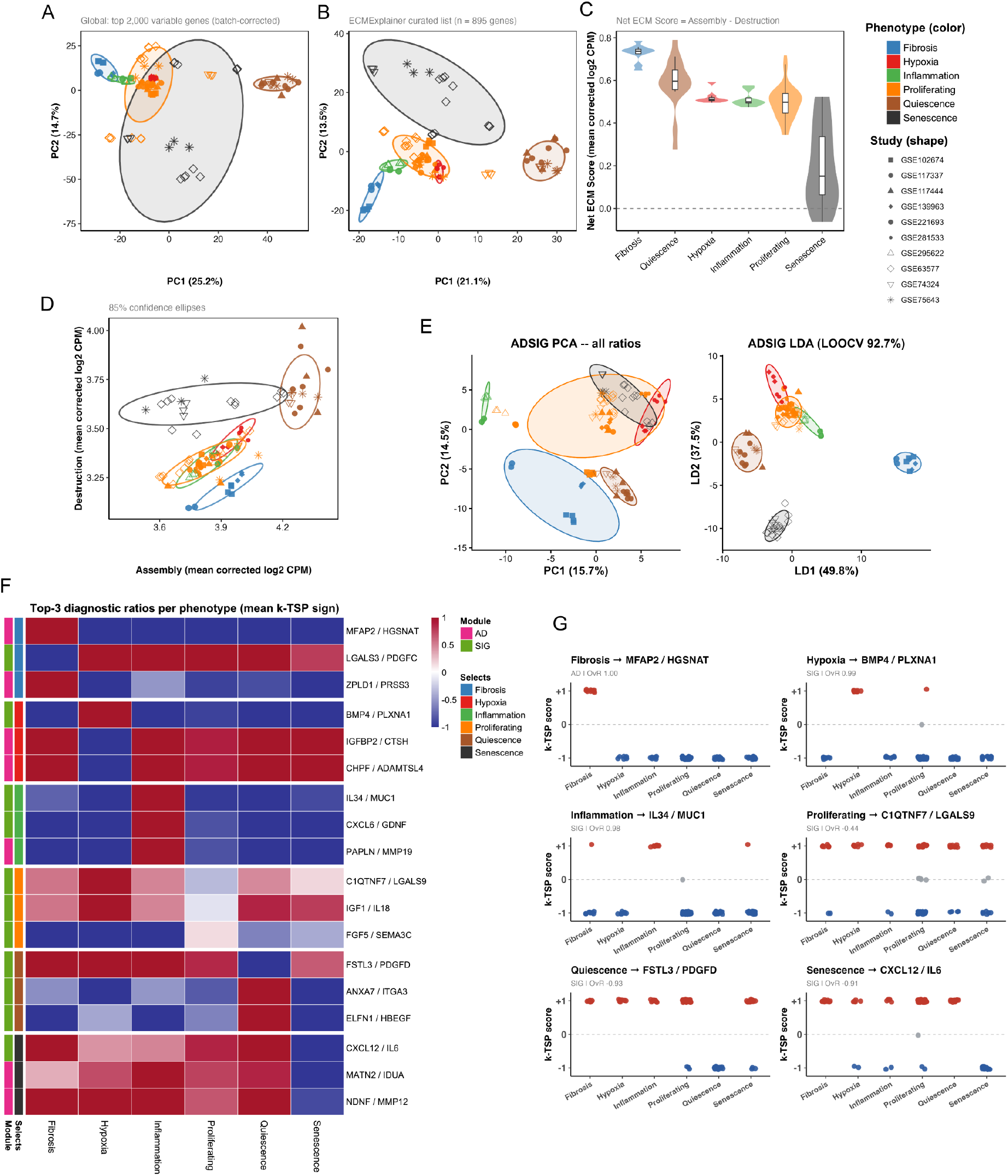
ECM transcriptional identity across fibroblast cell states. (A) Principal component analysis (PCA) of the 2,000 most variable genes genome-wide, following limma batch correction, showing global transcriptional separation of the six states. (B) PCA restricted to the ECMExplainer curated matrisome gene set (n = 895 genes retained after a variance filter of >0.005). (C) Net ECM Score per sample, defined as the mean batch-corrected log2 CPM of Assembly-module genes minus that of Destruction-module genes. (D) Assembly versus Destruction module scores per sample, resolving the two components underlying the Net ECM Score in (C). (E) PCA (left) and linear discriminant analysis (LDA, right) of all sign-encoded ADSIG-180 diagnostic ratios, colored by phenotype and shaped by source study, with 85% confidence ellipses. (F) Heatmap of the top three diagnostic ratios per phenotype (mean k-TSP sign), annotated by gene-module origin (AD, SIG) and phenotype selection; row color intensity reflects the mean signed log2 ratio (Δlog2) across samples for each phenotype. (G) Discrete k-TSP scores (+1 red, −1 blue, 0 grey; the sign of Δlog2 = log2 A − log2 B) for one representative top-ranked diagnostic ratio per phenotype, plotted across all six phenotypes; points are jittered to show density and the dashed line marks Δlog2 = 0. Module origin (AD/SIG) and one-versus-rest (OvR) discrimination score are annotated per panel.

To characterize how ECM programs differ between phenotypes, we first asked whether the overall balance between matrix-building and matrix-degrading activity could resolve the six states. We computed a Net ECM Score per sample (mean batch-corrected expression of Assembly genes minus that of Destruction genes) as a directed, exploratory summary of this balance (Figure 1C). Consistent with established biology, Fibrosis displayed the highest Assembly-dominant score, reflecting TGFβ-driven matrix deposition ^26^, while Quiescence ranked second, consistent with matrix preservation under growth arrest ^27^. Senescence scored lowest, reflecting the elevated matrix remodeling associated with the SASP ^8^. However, plotting Assembly against Destruction scores directly showed that Proliferating, Hypoxia, and Inflammation remained largely overlapping (Figure 1D). Thus, a single aggregate measure of Assembly–Destruction balance was insufficient to resolve all phenotypes, motivating incorporation of the Signaling module and a finer-grained, gene-pair representation of ECM state.

To capture the rewiring of ECM programs across phenotypes, we adopted a gene-pair log-ratio framework inspired by the Top-Scoring Pairs (TSP) algorithm ^28^ and its k-TSP extension ^29^. Rather than using absolute expression levels, this approach asks a simple question within each sample: which of the two genes is more highly expressed? For each gene pair (A and B), we compared log₂(CPM + 1) between the two genes and retained only the direction of that comparison: +1 when gene A was higher, -1 when gene B was higher, and 0 when the two were tied (Methods). Because both genes are measured in the same library and undergo the same normalization, any sample-wide scaling factor such as library size affects both genes equally and cancels in the comparison, so the classifier depends only on the relative ordering of the two genes within each sample. Candidate ratios were drawn from two biologically motivated pair classes. Assembly × Destruction (A/D) pairs encode the relative balance between ECM production and degradation, whereas Signaling × Signaling (SIG) pairs capture the relative state of ECM signaling pathways. Together these complementary pair classes allow the classifier to simultaneously assess both the structural remodeling state of the matrix and the cell–matrix communication state.

We constructed ECMExplainer using a stepwise feature-selection pipeline that first generated the biologically informed candidate gene pairs, then scored each pair for how well it separated one phenotype from the other five, and finally assembled a compact, non-redundant panel in which every phenotype is equally represented.

Scoring proceeded in two stages: First, candidate ratios were screened by η² (eta-squared), computed on their continuous log-ratio, log_2_(CPMₐ + 1) – log_2_(CPM_b_ + 1), which measures the fraction of variation in that log-ratio across the training samples that is attributable to phenotype, discarding ratios for which phenotype explained 10% or less of that variation. Second, surviving ratios were converted to their sign (-1, 0, +1) and ranked by TSP delta, defined as the largest one-versus-rest difference in the frequency with which the numerator ranked above the denominator. Each ratio was assigned to the phenotype yielding its maximum TSP delta, so that every candidate enters the panel as a marker of one specific state. The panel was then assembled in a single pass through all surviving ratios in descending TSP-delta order, subject to two constraints: each phenotype could contribute N/6 ratios (N = final panel size), and no gene could appear in more than one ratio. The latter constraint prevented the panel from collapsing onto a few highly ranked genes paired with arbitrary partners which would make the ECMExplainer vulnerable to the absence or poor quantification of an individual gene in validation datasets. As ranking is global, while quotas are per phenotype, the most discriminative ratios are taken first overall, resulting in a finished panel that is balanced by construction, with each phenotype contributing the same number of its own strongest ratios.

Starting from a total panel size of 60 ratios (10 ratios per phenotype), panel size was evaluated in increments of 30 ratios until the eligible ratio pool was exhausted. Classification was performed by projecting the sign-encoded ratio matrix into a low-dimensional PCA space, retaining the first ten principal components and then applying Linear Discriminant Analysis (LDA), a supervised method that finds the linear combination of input features maximally separating predefined classes. Accuracy at each panel size was assessed by both leave-one-out cross-validation (LOOCV) and the more stringent nested leave-one-study-out (LOSO) cross-validation, in which all samples from one study were withheld simultaneously to simulate deployment on an unseen dataset. The accuracy did not differ significantly between the different panel sizes tested (Supplementary Table 3). We therefore retained the 180-ratio panel because it provides 30 ratios per phenotype, spans 360 distinct genes, and is more tolerant of genes absent from external datasets than smaller panels while providing a broader representation of ECM gene space for future extension to additional phenotypes.

LOOCV at this 180-ratio panel size on the full 96-sample training set was 92.7% (macro-F1 = 0.92), with a LOSO of 47.9%. Residual misclassifications concentrated at the Proliferating–Hypoxia/Inflammation boundary. To further confirm suitability of this selected panel, we analysed a phenotype-level permutation comprising 2,000 permutations per ratio. 171 of 180 ratios (95.0%) showed greater one-versus-rest sign separation than expected by chance after Benjamini-Hochberg correction, confirming the ratio selection was successful. The nine ratios that did not meet this threshold all belonged to the Proliferating quota, consistent with Proliferating being distinguished by the absence of strong ECM shifts defining the other phenotypes. The final ECMExplainer classifier consists of sign encoding of the selected ratios followed by PCA and LDA, without any explicit batch-effect correction step (Figure 1E).

The 180 ratios comprising the ECMExplainer panel are unevenly distributed across the two functional modules: 76 ratios are Assembly/Destruction (A/D) pairs and 104 are Signaling/Signaling (SIG) pairs (Supplementary Table 3). This asymmetry is itself phenotype-specific: Fibrosis draws predominantly on A/D ratios (21/30), consistent with fibrotic identity being more distinctly defined by matrix turnover balance, whereas Proliferating and Hypoxia draw predominantly on SIG ratios (24/30 and 21/30, respectively), consistent with these states being more distinctly defined by receptor– ligand signaling co-expression than by matrix turnover balance. We generated heatmaps of the full 180-ratio panel, split by module, to visualize the mean sign of each ratio per phenotype alongside its hierarchical clustering (Supplementary Figure 3-4; Supplementary Table 3). In both the A/D and SIG heatmaps, we observe that phenotypes separate into distinct blocks of concordant ratio sign, and that the two module-specific dendrograms group phenotypes differently from one another. For example, the A/D dendrogram places Fibrosis as the most distinct outgroup, while the SIG dendrogram instead pairs Hypoxia and Inflammation as the closest pair. At single-sample resolution, hierarchical clustering of the full sign-encoded ratio matrix across all 96 samples (Supplementary Figure 5) showed that samples grouped predominantly by phenotype rather than by source study, with Fibrosis, Senescence and Quiescence forming the most internally coherent, study-independent blocks. This sample-level structure corroborates the phenotype-driven separation already evident in the LDA projection (Figure 1E) and demonstrates that the ratio-sign encoding preserves phenotype identity as a dominant axis of variation even before any supervised dimensionality reduction is applied.

### 3.2 ECMExplainer ratio panel encodes coordinated ECM program shifts

To understand how the selected ratios encode phenotypic identity, we asked, for each of the 180 ratios, how far each of its two constituent genes shifted in rank position between its selecting phenotype and Proliferating control samples from the same study (or, for ratios selecting Proliferating, between Proliferating and all other samples from the same study; Methods, Supplementary Table 3). Two features of each pair of shifts determine how the ratio works: whether the two genes move in the same or opposite direction, and how the total movement is divided between them.

To characterize the expression changes underlying the 180 gene ratios, we classified them into four patterns based on the direction and relative magnitude of changes in the two genes. In 72 ratios (40.0%), both genes shifted in the same direction but to different extents, so the ratio reports a change in the balance between two co-regulated genes (concordant shift’). The remaining 108 ratios showed changes in opposite directions. Among these, 44 ratios (24.4%) were dominated by changes in the numerator genes (Gene A dominant’), 28 (15.6%) by the denominator gene (Gene B dominant’), and 36 (20.0%) showed substantial contributions from both genes (reciprocal shift). Dominance was defined as one gene accounting for at least 70% of the combined absolute expression change. Detailed results for each ratio, including expression changes, confidence intervals, classification, and LD1-LD5 coefficients are given in Supplementary Table 3.

Having established the general behavior of the panel, we examined the leading discriminative ratios for each phenotype. The three highest-scoring one-versus-rest (OvR) ratios per phenotype are shown as their mean k-TSP sign across the cohort (Figure 1F), along with the single top ratio per phenotype at the sample level (Figure 1G) and shift level (Supplementary Figure 6). Fibrosis was identified with a perfect score by MFAP2/HGSNAT (A/D; OvR 1.00), a ‘Gene A dominant’ ratio in which the microfibril-associated glycoprotein MFAP2 rises, a direct readout of structural matrix deposition, consistent with MFAP2 upregulation in pulmonary and liver fibrosis ^30,31^. Hypoxia was distinguished by BMP4/PLXNA1 (SIG; OvR 0.99), again ‘Gene A dominant’, capturing induction of BMP4-mediated ECM signaling, consistent with hypoxic activation of BMP–SMAD pathways in cells ^32^. Quiescence was captured by FSTL3/PDGFD (SIG; OvR −0.93), a ‘Gene B dominant’ ratio in which PDGFD rises relative to a modest fall in FSTL3. PDGFD is known to play a role in cellular differentiation ^33^.

Inflammation was identified by IL34/MUC1 (SIG; OvR 0.98). Although both genes move in opposite directions reflecting coordinated elevation of the cytokine IL34 alongside suppression of MUC1 in the inflammatory ECM state ^34,35^, IL34 accounted for 81% of the movement, placing this as ‘Gene A dominant’.

Senescence provided the clearest reciprocal example in the panel with the two genes contributing almost equally. CXCL12/IL6 (SIG; OvR −0.91) showed CXCL12 falling while IL6 rose encoding the transition from homeostatic stromal chemokine activity to the pro-inflammatory SASP secretory program, with IL6 an established core SASP factor and loss of stromal CXCL12 a documented feature of senescent fibroblasts ^36,37^.

Proliferating fibroblasts were marked by C1QTNF7/LGALS9 (SIG; OvR −0.44), the weakest leading ratio, whose ratio-level shift was not distinguishable from zero (Supplementary Figure 6). This is consistent with their central, overlapping position in both the ECM PCA and ratio LDA. The Proliferating state is defined by the absence of strong ratio shifts that mark the other five phenotypes rather than by a distinctive ECM signature of its own.

Together these analyses show that the ECM phenotypes are encoded not by isolated changes in single transcripts, but by shifts in the relative ordering of functionally paired genes, with the balance of Assembly, Destruction and Signaling programs carrying the discriminative signal.

### 3.3 Validating the ECMExplainer on Senescence

To assess ECMExplainer’s performance on independent data, we first validated it against a well-characterized senescence dataset (SRP113329; Georgilis et al. 2018) comprising IMR90 ER:RAS^G12V^ fibroblasts sequenced six days after Ras-induced senescence (RIS) with 4-hydroxytamoxifen, alongside proliferating scrambled-siRNA controls (Proliferating). Within-dataset PCA confirmed clean separation of the two conditions (PC1 66.2%; Figure 2A) and projecting these samples into the ECMExplainer LDA space showed each sample positioned closest to its true phenotype (Figure 2B). We quantified phenotype assignment using the LDA posterior probabilities. For each sample, the model assigns probabilities across the six training phenotypes, which sum to one, and the phenotype with the highest posterior probability is reported as prediction (Figure 2C). We additionally calculated each sample’s Euclidean distance to the centroid of each ECMExplainer training phenotype in the LDA space, which we refer to hereafter as a distance heatmap (Figure 2D).

**Figure 2:**
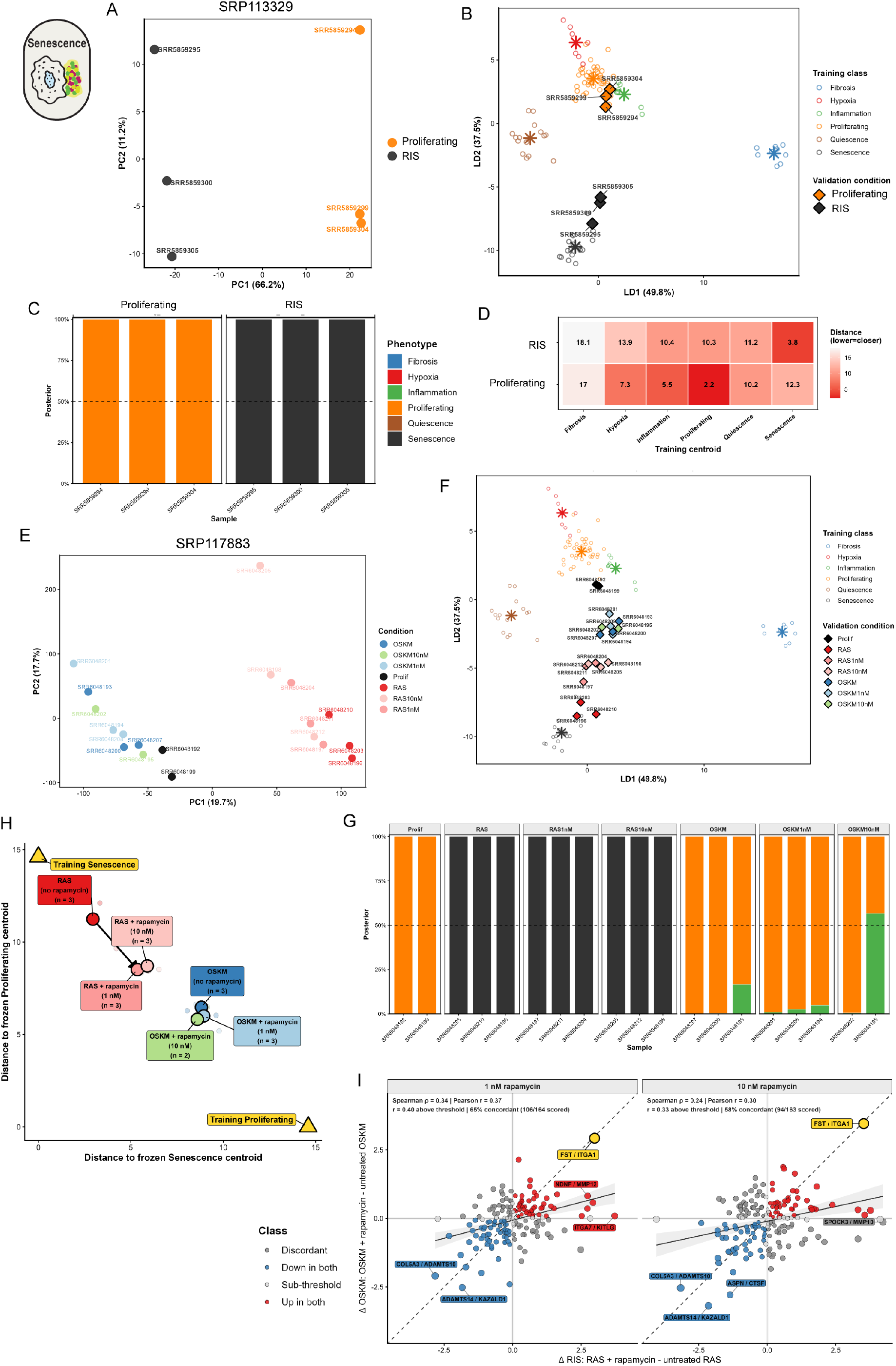
ECMExplainer detects Ras-induced senescence and rapamycin-mediated ECM rescue. (A) PCA of SRP113329. (B) Two-dimensional LDA projection of SRP113329 samples on the ECMExplainer training phenotypes. (C) Stacked posterior probability bars for each SRP113329 sample. Dashed line marks the 50% confidence threshold. (D) Median Euclidean distance heatmap from each SRP113329 condition to each ECMExplainer training phenotype. (E) PCA of SRP117883. (F) Two-dimensional LDA projection of SRP117883 samples on the ECMExplainer training phenotypes. (G) Stacked posterior probability bars for each SRP117883 sample as in (C). (H) ECM-state response to rapamycin in RAS-and OSKM-induced senescent IMR90 fibroblasts, plotted as Euclidean distance to the frozen Senescence versus Proliferating training centroids in ECMExplainer LDA space. Faint points are individual samples; large outlined points are per-condition mediods labeled with treatment, dose and n. Solid arrows trace RAS (untreated → 1 nM → 10 nM rapamycin), dashed arrows trace OSKM. Gold triangles mark the training centroids. Axes are measured distances; arrows interpolate between observed doses. Net repositioning is quantified by D_Prolif − D_Sen (see Methods). (I) Rapamycin-induced ECM-ratio shifts in RAS versus OSKM, faceted by dose. Each point is one of the 180 k-TSP ratios: x-axis, RAS + rapamycin minus untreated RAS; y-axis, OSKM + rapamycin minus untreated OSKM (log_2_). Colors denote concordance class: "Up in both" and "Down in both" (concordant), "Discordant," and "Sub-threshold" (|Δ| ≤ 0.05 log₂ in either arm, shaded band, excluded from concordance scoring). FST/ITGA1 is gold. The black line is the ordinary least squares linear fit with 95% CI. In panel text reports Spearman ρ, Pearson r (all ratios and above threshold subset), and percent concordant among scored ratios.

Next, we analyzed an IMR90 fibroblast dataset containing Ras-induced senecence (RAS) and OSKM-associated reprogramming-induced senescence (OSKM), each with untreated, 1 nM rapamycin-treated, and 10 nM rapamycin-treated conditions, allowing us to test the ECMExplainer on classical and specialized modes of senescence (Figure 2E) (SRP117883, Aarts et al. 2017). As we saw in SRP113329, both the untreated (Prolif) and RAS samples were correctly predicted and mapped to the expected centroids (Figure 2F). In contrast, OSKM samples were classified closer to Proliferating than Senescence, in line with their position between the Proliferating and Senescence centroids in the LDA space (Figure 2F) and with the original within-dataset PCA, where OSKM samples clustered closer to proliferating controls than to RAS-senescent cells (Figure 2E). This is consistent with the known transcriptional ambiguity of reprogramming-associated senescence, which shares features with both a proliferative intermediate and a classical senescent state ^39^.

Rapamycin shifted RAS samples away from the Senescence centroid and toward the Proliferating centroid irrespective of rapamycin dose (Figure 2G-H), consistent with rapamycin’s established senomorphic activity, partially reversing the senescent ECM program without fully converting it to a proliferative state ^40^. The same treatment produced little directional movement of OSKM samples, which remained at their intermediate Proliferating-classified location regardless of dose (Figure 2H).

We therefore computed the rapamycin-induced shifts within the two models: RAS plus rapamycin minus untreated RAS, versus OSKM plus rapamycin minus untreated OSKM. Across the 180 ratios, the shift patterns were partially concordant at both rapamycin doses (Figure 2I). The single most concordantly upregulated ratio in both conditions was FST/ITGA1 (Follistatin/Integrin α1; Δ_RIS ≈ +3.52, Δ_OSKM ≈ +3.45 at 10 nM), one of the Senescence-selected ECMExplainer ratios from the training set (Signaling module; TSP δ = 0.78; direction: down in Senescence), demonstrating that rapamycin modulates a feature of the ECM senescence signature even in a model where the overall phenotype classification does not change.

We next applied ECMExplainer to a replicative senescence dataset (SRP104418) comprising human cardiac fibroblasts ^41^. The dataset includes untreated early passage cells (EarlyPass, population doubling [PD] 30), untreated late passage cells (LatePass, PD52), and cells cultured under rapamycin treatment (RapaCtrl), methionine restriction (MetR), or combined rapamycin plus methionine restriction (RapaMetR) at PD45. The combined RapaMetR trajectory was also examined at late passage (OldRapaMR, PD85) (Figure 3A).

**Figure 3:**
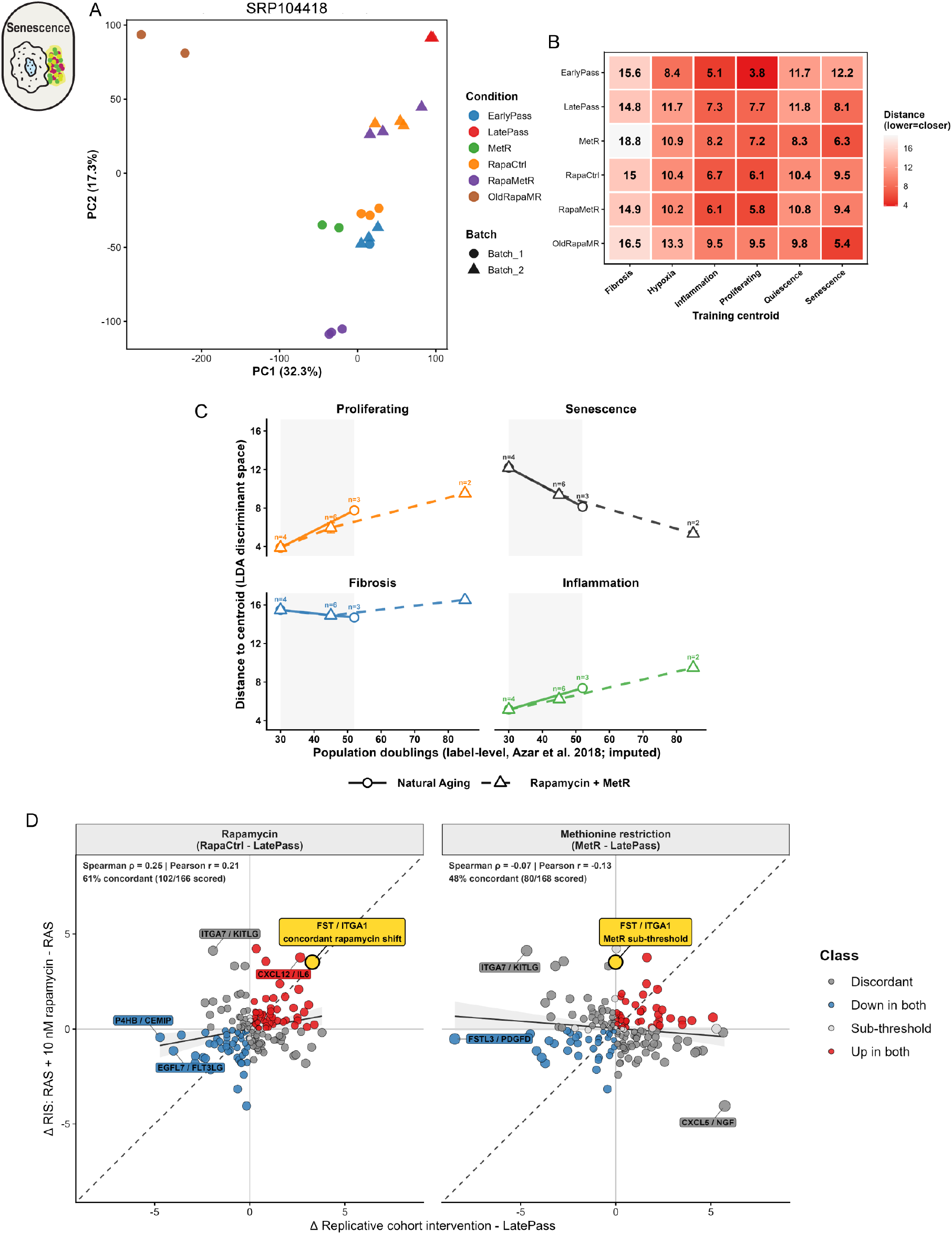
ECMExplainer tracks replicative matrisome drift and identifies FST/ITGA1 as a cross-context rapamycin-responsive biomarker (A). PCA of SRP104418. (B) Median Euclidean distance heatmap from each SRP104418 condition to each ECMExplainer training phenotype. (C) Four-panel matrisome-drift kinetics. Mean Euclidean distance (± SE; n per knot indicated) from projected human cardiac fibroblast samples to the frozen Proliferating, Senescence, Fibrosis, and Inflammation ECMExplainer centroids, in LDA discriminant space, plotted against population doubling. Solid lines/circles, Natural Aging arm (EarlyPass PD 30 → LatePass PD 52); dashed lines/triangles, Combined (Rapamycin + MetR) arm (EarlyPass PD 30 → RapaCtrl PD 45 → OldRapaMR PD 85). Shaded band (PD 30–52) marks the overlapping support where the rate comparison is valid; connecting lines are visual guides, not fitted kinetics. (D) Cross-study conservation of the rapamycin ECM-ratio signature. Each point is one of the 180 shared ADSIG-180 k-TSP ratios, plotted as its Ras-induced shift (Δ_RIS = RAS + 10 nM rapamycin − RAS, SRP117883; y-axis) against its replicative-cohort intervention shift (SRP104418; x-axis), colored by four-way concordance class ("Up in both," "Down in both," "Discordant," "Sub-threshold" = |Δ| ≤ 0.05 log₂ in either axis, shaded band, excluded from scoring). Left, rapamycin (RapaCtrl − LatePass); right, methionine-restriction control (MetR − LatePass). Facet headers report Spearman ρ, Pearson r, and percent-concordant among scored ratios. The black line is the OLS fit with 95% CI; labelled ratios are the top 5 concordant and top 3 discordant by |Δ| sum.

ECMExplainer correctly classified EarlyPass cells as Proliferating (mean distance to the Proliferating centroid = 3.9), while LatePass cells moved toward the Senescence pole (distance falling from 12.2 at PD30 to 8.1 at PD52) (Figure 3B). In the intervention arms at PD45, RapaCtrl and RapaMetR remained closer to the Proliferating centroid than MetR alone which was closer to the Senescence centroid. This indicated that rapamycin, not methionine restriction, conferred protection against ECM senescence. This protection though was PD-dependent. OldRapaMR (the combined RapaMetR cells at PD85) had drifted back toward the Senescence centroid, demonstrating that combined intervention delays but does not permanently prevent ECM replicative senescence progression (Figure 3C).

To test whether the ECM signature associated with rapamycin was shared across distinct senescence contexts, we compared the only directly comparable single-agent intervention available in both datasets: rapamycin treatment in replicatively aged cardiac fibroblasts (RapaCtrl – LatePass, SRP104418) and in Ras-induced senescent IMR90 fibroblasts (RAS + 10 nM rapamycin, SRP117883). Methionine restriction was also included as a non-rapamycin comparator within SRP104418, and is not a matched treatment across the two datasets.

Across the 180 ECMExplainer ratios, the rapamycin shifts were modestly but positively correlated between replicative and Ras-induced senescence (Spearman ρ = 0.25, Pearson r = 0.21; 61% of scored ratios concordant, Figure 3D, left). FST/ITGA1 (Follistatin/Integrin α1) was the most concordant ratio increasing in both models (Δ_Replicative = +3.29, Δ_RIS = +3.52; combined magnitude 6.81). FST/ITGA1 is a Senescence-associated ratio in the ECMExplainer panel, with the denominator ITGA1 preferentially ranked above FST in Senescence (Supplementary Table 3).

This cross-model concordance was not reproduced by methionine restriction. Substituting MetR-LatePass for the replicative rapamycin contrast eliminated the association with the Ras-induced rapamycin response (ρ = −0.07, r = −0.13), and FST/ITGA1 showed no detectable shift under MetR (Δ_MetR = −0.01, sub-threshold, Figure 3D, right). That the same ECM ratio moves in the same direction under two different senescence triggers, but only in response to rapamycin, not methionine restriction, indicates a drug-specific, conserved matrisome axis rather than generic cross-study correlation.

### 3.4 Validating the ECMExplainer on Fibrosis

To validate Fibrosis, we chose a dataset that contained primary cardiac fibroblasts derived from patients undergoing open-heart surgery, that were cultured *in vitro*, and either left untreated or treated with TGF-β1 to induce fibrosis (SRP102952, Schafer et al. 2017). In a global PCA, the two conditions were not resolved by the leading principal components (Figure 4A), reflecting the inter-patient heterogeneity expected of primary cells from 84 individual donors. Instead, the samples separated more clearly on PC4 (4.5% of total variance), confirming that the TGF-β1 response is recoverable. In contrast, projection into the ECMExplainer LDA space revealed a clear shift: median distance to the Fibrosis centroid decreased from 7.6 (Baseline) to 5.9 (TGF-β1-treated), while distance to all other centroids increased (Figure 4B). Nevertheless, many Baseline individual samples were predicted as Fibrosis. We attribute this to the fibroblasts being derived from older patients with end-stage failing hearts. Importantly, when we looked per patient, we could see a general shift from the Proliferating to Fibrosis centroid on TGF-β1 treatment (Figure 4C) with 95.2% of patients showing the shift toward Fibrosis (paired Wilcoxon signed-rank p = 2×10⁻¹⁴; Figure 4D), confirming the ECMExplainer’s sensitivity to detect fibrotic ECM remodeling even against a potentially high-fibrosis baseline.

**Figure 4:**
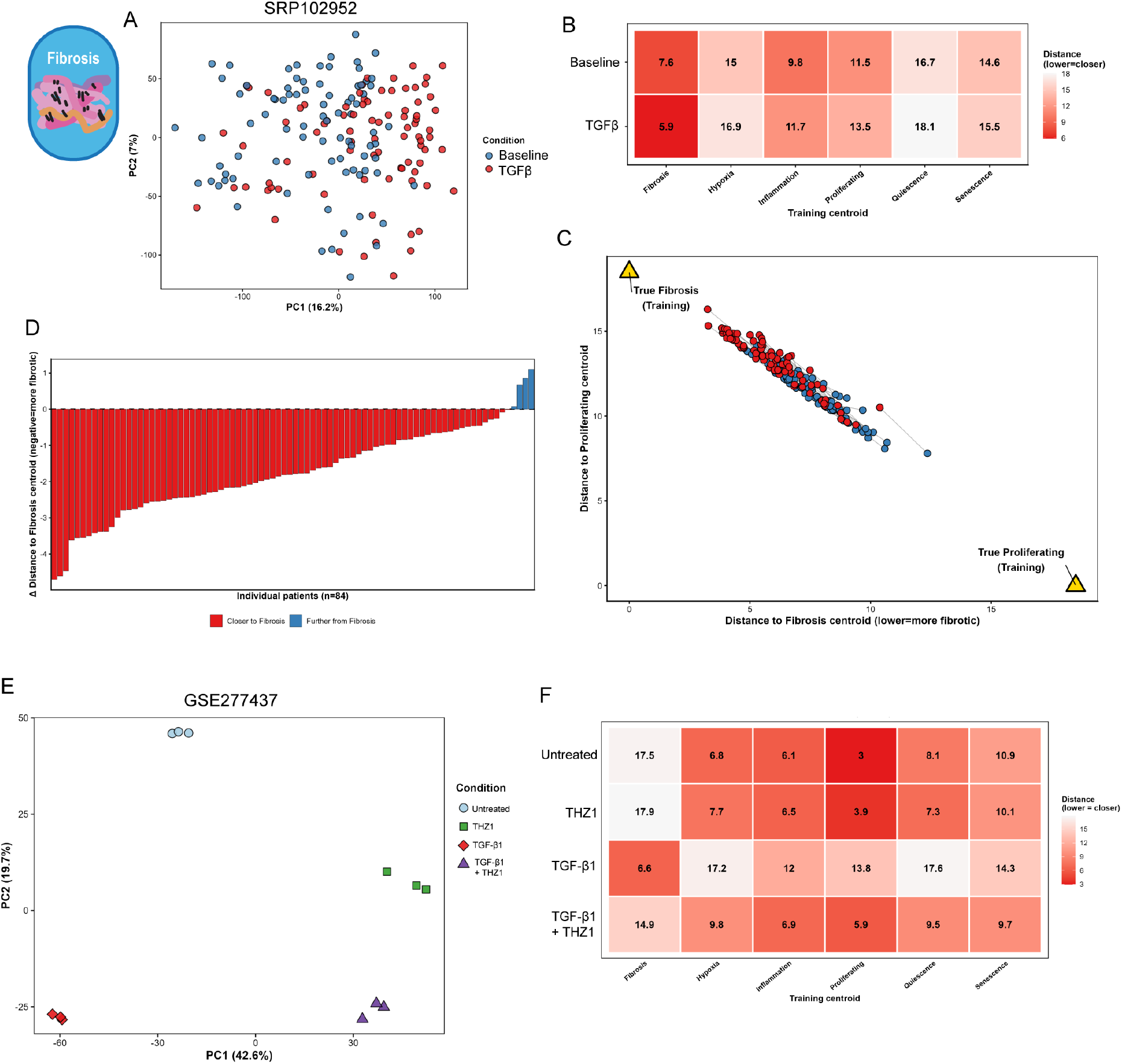
Validating ECMExplainer on two fibrosis datasets: (A) PCA of SRP102952 validation samples. (B) Median Euclidean distance heatmap from each SRP102952 sample to each ECMExplainer phenotype. (C) Two-dimensional paired patient trajectory plot in Fibrosis-Proliferating space. Gold triangles mark the positions of the true training centroids. (D) Patient-level waterfall plot of the change in Euclidean distance to the Fibrosis centroid after TGF-β1 stimulation. (E) PCA of GSE277437 validation samples. (F) Median Euclidean distance heatmap for GSE277437.

We further validated fibrosis detection using an independent lung fibroblast dataset (GSE277437; Ma et al. 2025), chosen because it tests both the induction and the pharmacological prevention of a fibrotic matrix state. Primary human lung fibroblasts (Untreated) were differentiated into activated myofibroblasts by TGF-β1 treatment, with parallel arms receiving the CDK7/12/13 inhibitor THZ1 alone or THZ1 co-administered with TGF-β1. Global PCA separated all four arms (Figure 4E), while ECMExplainer correctly classified all TGF-β1-treated samples as Fibrosis and all Untreated and THZ1-alone samples as Proliferating. Critically, co-treatment with THZ1 largely prevented the TGF-β1-driven displacement, with co-treated samples being predicted as Proliferating (Figure 4F). Expressed on the axis spanning the TGF-β1 and untreated states, this corresponded to a prevention of 76% of the fibrotic ECM shift. However, we can see that the co-treated cells were not globally restored since they remain displaced from the other conditions in the global PCA (Figure 4E). ECMExplainer therefore resolved the specific reversal of the matrix phenotype from the drug’s global transcriptional effect, demonstrating that the classifier tracks a pharmacologically modifiable ECM state rather than treatment identity per se.

### 3.5 Additional Validation Sets

To further assess the generalizability of ECMExplainer, we analyzed thirteen additional independent publicly available fibroblast transcriptome datasets encompassing four phenotypes (Senescence, Quiescence, Proliferating and Fibrosis, Supplementary Figures 7–10). Across five senescence datasets, including three replicative (SRP121031, ^44^; SRP034541, ^45^; SRP040745, ^46^), one 10-day stress-induced (SRP098713, ^47^), and one time-course stress-induced (SRP136071, ^48^), senescent samples showed consistently shorter median distances to the Senescence centroid than their paired Proliferating controls across all five datasets (Supplementary Figure 7). SRP136071 further revealed that the ECMExplainer captures the progressive maturation of the senescent matrisome (Supplementary Figure 7A). Within the first two days of induction, samples remained close to the Proliferating centroid (1.9-3.2, versus 1.6 in proliferating controls). A clear shift emerged only by day 4, where distance from the Proliferating centroid reached its maximum of 5.3 and distance to the Senescence centroid fell to its minimum of 8.4. After, both measures stabilized at days 6 and 8. The ECM program therefore does not shift immediately but is acquired over several days before stabilizing, consistent with the known temporal acquisition of the senescent secretome and matrisome ^49,50^.

In two further datasets containing both senescence and quiescence conditions (SRP017142, ^51^; ERP021140, ^52^), senescent and quiescent samples moved closer to their expected centroids and away from Proliferating (Supplementary Figure 8). Across four quiescence-only datasets (SRP153724, ^53^; SRP153205,^54^; SRP052706, ^55^; SRP045867, ^56^), contact-inhibited or serum-starved fibroblasts were classified closest to the Quiescence centroid (median Dist_Quiescence 3.6–6.0), and paired Proliferating controls to Proliferating (median Dist_Proliferating 1.8–3.6), with 15 of 16 samples correctly classified (Supplementary Figure 9).

Finally, the Fibrosis axis was validated on two independent TGF-β1-stimulated fibroblast datasets (GSE302114, ^57^; GSE273882, ^58^). In both, untreated fibroblasts localized closest to the Proliferating centroid (Dist_Proliferating 3.3 and 3.9 respectively), while TGF-β1-treated cells were correctly classified as Fibrosis (Dist_Fibrosis 7.0 and 6.9 respectively), confirming that the ECMExplainer’s fibrotic matrisome signature generalizes across lung and other fibroblast lineages (Supplementary Figure 10).

Together, these thirteen independent datasets further establish that the ECMExplainer correctly distinguishes all four ECM states (Proliferating, Senescence, Quiescence, and Fibrosis), whether through correct prediction, or via analysis of the centroid distances, consistent with its design as a cross-context matrisome classifier.

### 3.6 The ECM Aging Trajectory Diverges between chronological and Progeroid Aging

Having validated the ECMExplainer on distinct biological scenarios, we next applied it to chronological aging in human dermal fibroblasts. The publicly available RNA-sequencing dataset GSE113957 comprises cultured dermal fibroblasts established from skin biopsies of 133 healthy human donors aged 1–94 years and 10 patients with Hutchinson-Gilford Progeria Syndrome (HGPS), a premature aging disorder ^59^. Healthy donors were binned into four age groups: Young (≤20 years, n=26), Adult (20– 45 years, n=38), Middle (45–70 years, n=32), and Aged (>70 years, n=37), with HGPS samples treated as an accelerated aging comparator (Figure 5A).

**Figure 5:**
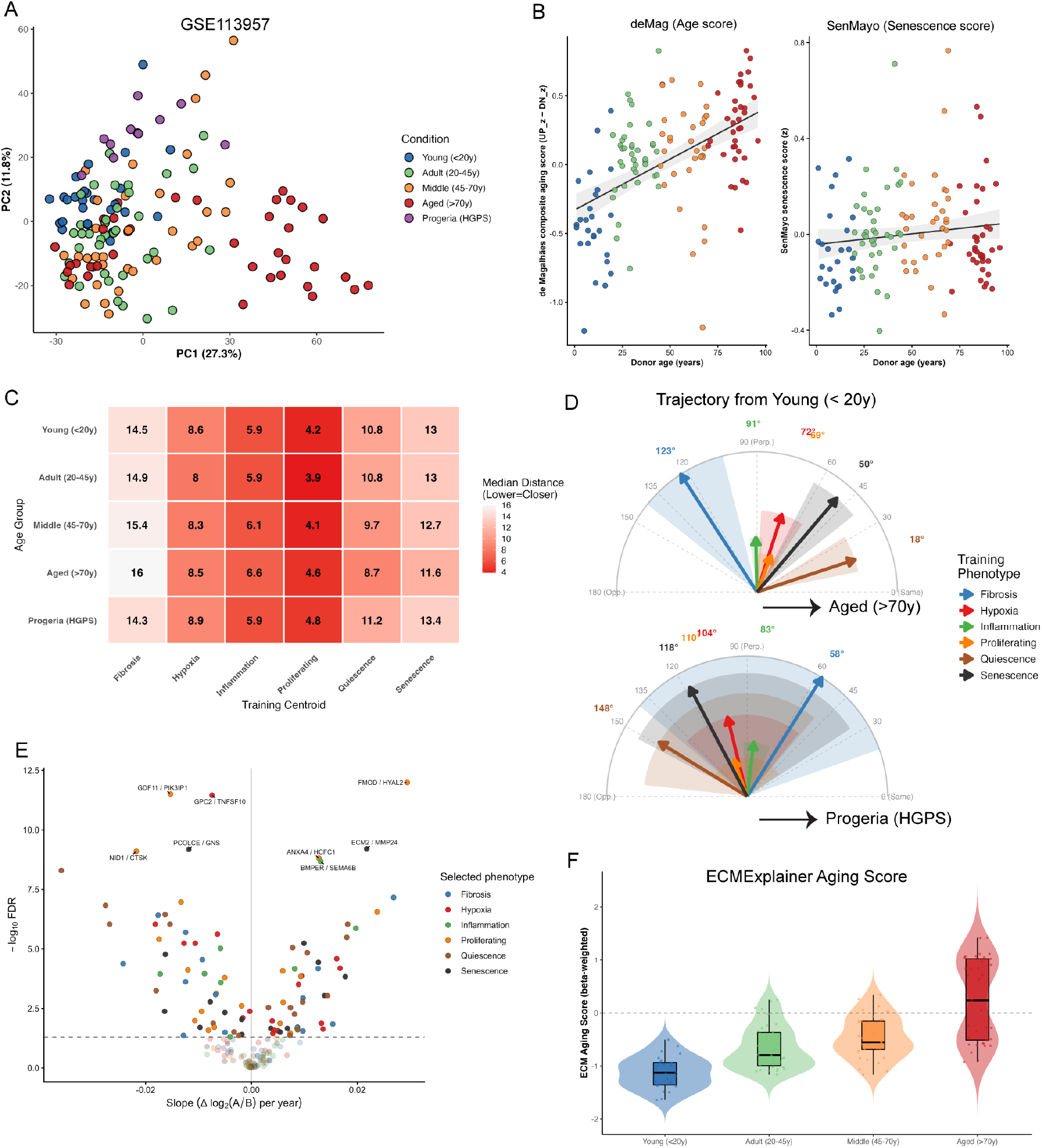
ECMExplainer characterizes human chronological aging. (A) PCA for all samples in GSE113957 (n=133 healthy donors, 1–94 years; n=10 HGPS). (B) Sample-level scores for the de Magalhães composite aging gene signature (left; ^60^) and the SenMayo senescence gene set (right;^61^), plotted against donor age. Lines show linear regression fit with 95% confidence intervals. (C) Median Euclidean distance heatmap from each GSE113957 age group to each training phenotype centroid. (D) Trajectory protractor plots quantifying the angular alignment between each ECMExplainer training phenotype vector (colored arrows) and the reference biological trajectory (grey arrow at 0°): Young <20y → Aged >70y (left) and Young <20y → HGPS (right). Phenotype arrows originate at the Young centroid; arrow length is proportional to Euclidean distance from the Young centroid in LDA space, normalized to the most distant phenotype. The shaded wedge around each phenotype arrow denotes the bootstrap 95% confidence interval of the trajectory angle, estimated by independently resampling the Young and Aged/HGPS groups with replacement (n = 2,000 iterations). Angles closer to 0° indicate stronger alignment with the trajectory; 90° = orthogonal; 180° = opposite direction. (E) Volcano plot of per-ratio linear regression slopes (Δ log₂(A/B) per year) against −log₁₀ FDR across the 133 healthy donors, colored by the ECMExplainer training phenotype that selected each ratio. Dashed horizontal line: FDR = 0.05. Selected top ratios are labelled. (F) Sign-based ECM aging score per chronological age bin.

We first confirmed that the cohort retained a chronological-aging transcriptional signal. A signed composite score based on the de Magalhães (deMag) composite aging gene signature, derived from a cross-species meta-analysis of 27 transcriptomic datasets ^60^, correlated with donor age among healthy individuals (Spearman ρ = 0.56, p = 1.5×10⁻¹², Figure 5B). In contrast, the SenMayo senescence gene set, a validated 125-gene panel for identifying senescent cells in vivo, was not associated with age (ρ = 0.10, p = 0.25; Figure 5B, Saul et al. 2022). Together, these results suggest that the transcriptional changes observed with age primarily represent a general aging program rather than increased cellular senescence.

ECMExplainer assigned 141 of 143 samples to the Proliferating class (Supplementary Figure 11A), consistent with their shared expansion as dermal fibroblast cultures. Despite this common discrete label, median centroid distances showed a graded age-associated shift toward the Senescence centroid (13.0 → 13.0 → 12.7 → 11.6) and the Quiescence centroid (10.8 → 10.8 → 9.7 → 8.7) (Figure 5C). These continuous centroid-distance changes indicate that chronological aging modifies the ECM-ratio profile even when the nearest-centroid class remains unchanged.

To compare the direction of these shifts we developed a polar visualization of vector-angle analysis in ECMExplainer discriminant space which we term a ‘protractor plot’. The Young-to-Aged centroid vector was defined as the 0° reference, and angles between this vector and Young-to-training-phenotype vectors were computed in the full five-dimensional LDA space, with bootstrap 95% confidence intervals (n=2,000 iterations; Figure 5D). The corresponding centroid/vector geometry is shown in Supplementary Figure 11B. Quiescence aligned most closely with the chronological aging vector (18°, 95% CI 11–32°), followed by Senescence (50°, 95% CI 38–63°), consistent with the known accumulation of both non-proliferative ECM states in aged tissue ^62,63^. In contrast, the Young-to-HGPS trajectory followed a distinct trajectory. Fibrosis aligned most closely with HGPS (58°, 95% CI 20–140°), while Inflammation was the next closest (83°, 95% CI 65–94°). Quiescence, on the other hand was near-opposite (148°, 95% CI 49–174°), indicating that progeroid accelerated aging drives an ECM program enriched for fibrotic and inflammatory remodeling rather than the quiescence-senescence axis of normal aging, consistent with the systemic fibrotic phenotype described in HGPS (20798379, 39273272).

We next tested each of the ECMExplainer ratios for association with chronological age across the 133 healthy donor-derived fibroblast lines, using linear regression of continuous log_2_-ratio values on age and Benjamini–Hochberg correction for multiple testing (Figure 5E, Supplementary Data Table 3). Ninety-six ratios (53.3%; 60 SIG, 36 AD) were age-associated at FDR < 0.05 (Supplementary Table 4). Of these, 47 increased and 49 decreased with age. The age-associated ratios were distributed across all six phenotype-defined subsets. Proliferating and Quiescence contributed the largest numbers (20 each), followed by Hypoxia (17), Senescence (16), Fibrosis (14) and Inflammation (9) (Figure 5E). Thus, the aging-associated matrisome program incorporates components of multiple phenotypic ECM states rather than being defined by a single state such as Senescence.

FMOD/HYAL2 showed the strongest association with age (slope = +0.029 log₂ units/year, FDR = 1.0×10⁻¹²; Proliferation-selected ratio), reflecting progressive upregulation of the extracellular matrix proteoglycan Fibromodulin (FMOD) relative to the hyaluronan-degrading enzyme HYAL2 (Supplementary Figure 11C). Both genes individually tracked age (FMOD: slope = +0.020/year, HYAL2: slope = -0.009/year), with their combined ratio capturing a more consistent age trend than either gene alone.

We then used the 96 age-associated ratios to derive a continuous ECM Aging Score. For each ratio, its age-regression slope was used as a weight. Then, because ECMExplainer operates on binary ratio signs, we also constructed a deployable sign-based version by replacing each continuous ratio with its direction only: +1 when Gene A > Gene B and -1 when Gene A < Gene B, while retaining the same age-derived weights. In GSE113957, the continuous and sign-based scores were highly correlated (r = 0.963), indicating that, within this cohort, ratio direction retained most of the age-related ordering captured by the continuous ratio magnitude. The sign-based ECM Aging Score increased across age bins (median: −0.36 Young, −0.22 Adult, −0.13 Middle, +0.09 Aged; Figure 5F), providing a single interpretable matrisome readout of biological age. Together, these analyses show that chronological aging follows a distinct ECM remodeling trajectory that can be summarized by a quantitative ECM Aging Score.

### 3.7 The ECM Aging Score Stratifies Successful From Failed OSKM-Reprogramming

Partial OSKM cellular reprogramming has been proposed as a route to reverse biological age, with rejuvenation demonstrated via DNA methylation clocks ^64,65^. Lu et al. (2025) recently described mesenchymal drift (MD), characterized by age-associated upregulation of mesenchymal/EMT-related genes across multiple human tissues, and reported that a short OSKM treatment suppresses this transcriptional program in aged fibroblasts. Although the biological interpretation of MD suppression in intrinsically mesenchymal cells remains unclear, of the 205 MD genes, 117 (57.1%) are present in the ECMExplainer master gene list (76 Assembly, 34 Signaling, 7 Destruction), and 48 appear directly in the 180-ratio panel, positioning the ECMExplainer to ask whether reprogramming success is legible at the level of the matrisome.

We applied the ECMExplainer to a longitudinal RNA-sequencing dataset (GSE165176, ^67^) profiling dermal fibroblasts from six donors: two young (Y1, Y2), two normal-age (N2, N3) and two old (O1, O2), undergoing Sendai virus-mediated OSKM reprogramming. At multiple timepoints between Day 0 (Baseline) and Day 54, cells were FACS-sorted into successfully reprogramming (SSEA4+) and failing-to-reprogram (CD13+) populations, yielding 124 RNA-sequencing samples (Figure 6A). Projected into the ECMExplainer LDA space, baseline fibroblasts mapped near the Proliferating centroid (Figure 6B). Both the SSEA4^+^ (reprogramming) and CD13^+^ (failing) populations initially moved away from Proliferating centroid over the first two weeks, consistent with the known early disruption of fibroblast identity, before diverging (Figure 6C). Between days 29 and 47, successfully reprogramming cells moved progressively away from both the Quiescence (8.7 → 11.4) and Senescence (8.3 → 11.8) centroids, indicating ECM de-differentiation away from age-associated matrix states. In contrast, failing cells drifted steadily closer to both Quiescence and Senescence between day 0 and day 21 (Proliferating distance: 3.6 → 7.3; Senescence distance: 12.6 → 7.9), consistent with failed reprogramming cells adopting an aged, non-proliferative ECM state.

**Figure 6:**
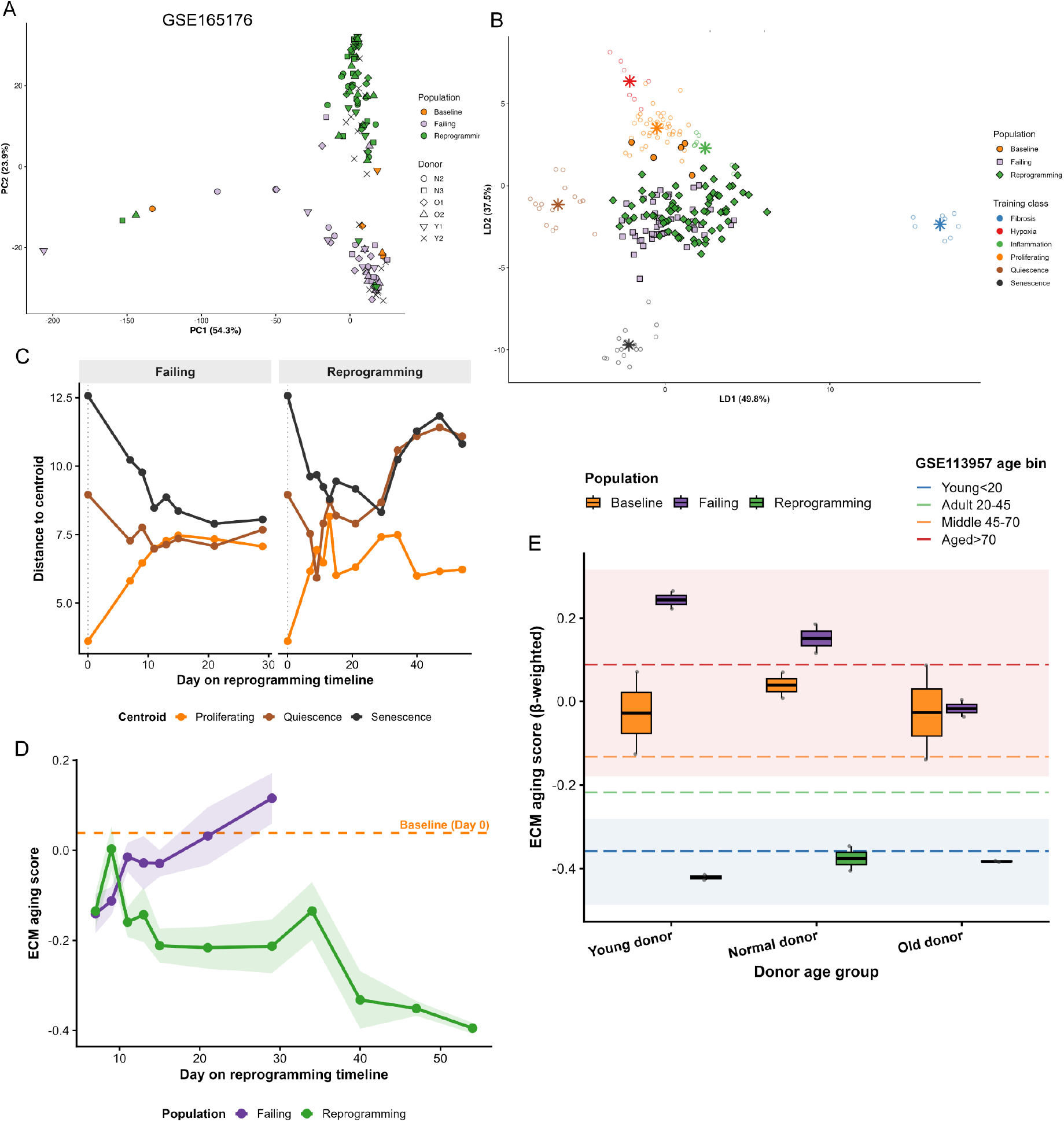
ECMExplainer applied to OSKM reprogramming of human fibroblasts (GSE165176, Gill et al., 2022). (A) Unsupervised PCA of log_2_RPM expression values across the 2,000 most variable genes. (B) Supervised LDA projection of the ECMExplainer ratio panel. Validation samples are overlaid. (C) ECM state trajectory. Median Euclidean distance from the ECMExplainer LDA centroids for Proliferating, Quiescence, and Senescence plotted over the reprogramming timeline for SSEA4+ reprogramming and CD13+ failing populations. (D) Sign-based ECM Aging Score over the reprogramming timeline for Failing (purple) and Reprogramming (green) populations. Lines show median per timepoint; shaded bands indicate interquartile range. Orange dashed line indicates the Baseline (Day 0) median. (E) Sign-Based ECM Aging Score at three biological endpoints, stratified by donor age group (n = 2 donors per age group). Baseline: single timepoint, Day 0 per donor. Failing endpoint: each donor’s final timepoint. Reprogramming endpoint: each donor’s final timepoint (day 54). Horizontal lines and shaded bands indicate the median ± interquartile range of each age bin (Young ≤ 20 y; Adult 20–45 y; Middle 45–70 y; Old > 70 y) from the GSE113957 reference cohort.

To quantify these trajectories in terms of biological age, we applied the sign-based ECM Aging Score, using the age-trend weights derived from GSE113957 without any refitting, to the reprogramming timeline in GSE165176 (Figure 6D). Stratified by donor age group at three biological endpoints (Baseline Day 0, Failing and Reprogramming at each donor’s final timepoint, day 21-29 and day 54 respectively) and overlaid against the GSE113957 age-bin reference bands for biological context only (Figure 6E), a striking divergence emerged. All six donors converged to ECM Aging Scores within or below the Young reference band (donor median −0.39, range −0.43 to −0.35) regardless of donor age. This reduction relative to the GSE165176 Baseline (Day 0) distribution was statistically significant by one-sided Wilcoxon rank-sum test (n=71, W=59, BH-adjusted p=0.0035, testing for a younger ECM signature). In contrast, failing donors drifted upward (scores −0.04 to 0.27, approaching the Aged band median of 0.09), though this drift was not significantly different from Baseline (n=47, W=115, BH-adjusted p=0.242).

Together, these data demonstrate that the ECMExplainer’s ECM Aging Score separates successful from failed OSKM reprogramming endpoints in this six-donor cohort, and that successful reprogramming drives convergence to a Young-equivalent ECM age irrespective of the donor’s chronological age. This suggests that matrisome expression alone may track reprogramming success without requiring methylation clocks or transcriptome-wide measures.

### 3.8 Benchmarking the ECMExplainer Against Competing Classifiers

In the preceding validation studies, we demonstrated that ECMExplainer correctly recovers sample phenotypes on independent datasets and quantifies geometric proximity to trained phenotype centroids. To determine whether this performance depends on the biologically informed selection of ECM gene pairs, rather than simply use of ECM genes or of ratio features in general, we benchmarked it against four control classifiers that include both axes of its design: feature selection and feature representation.

We first asked whether classification performance benefits from restricting gene selection to the ECM and pairing genes according to their biological functions. POOL-180 replaces the biologically distinct Assembly/Destruction and Signaling/Signaling pairings with unstructured ratios drawn from any ECM gene pair, isolating the contribution of pairing genes by biological function rather than at random. ALLGENE-180 removes the ECM restriction itself, selecting ratios by discriminative performance from a genome-wide, expression filtered gene universe. Both panels were reported at 180 ratios, such that the only variable between the three ratio panels is the gene universe and the pairing rule. Two further controls, All-Genes (limma) and ECM-genes (limma), replace ratios with absolute log-CPM expression, drawn from the full expressed transcriptome and the curated ECM gene set respectively, with limma batch correction applied at training time. All five models share the same downstream PCA→LDA architecture and are projected onto validation cohorts using identical reference-projection procedures with no retraining or leakage (Figure 7A).

**Figure 7:**
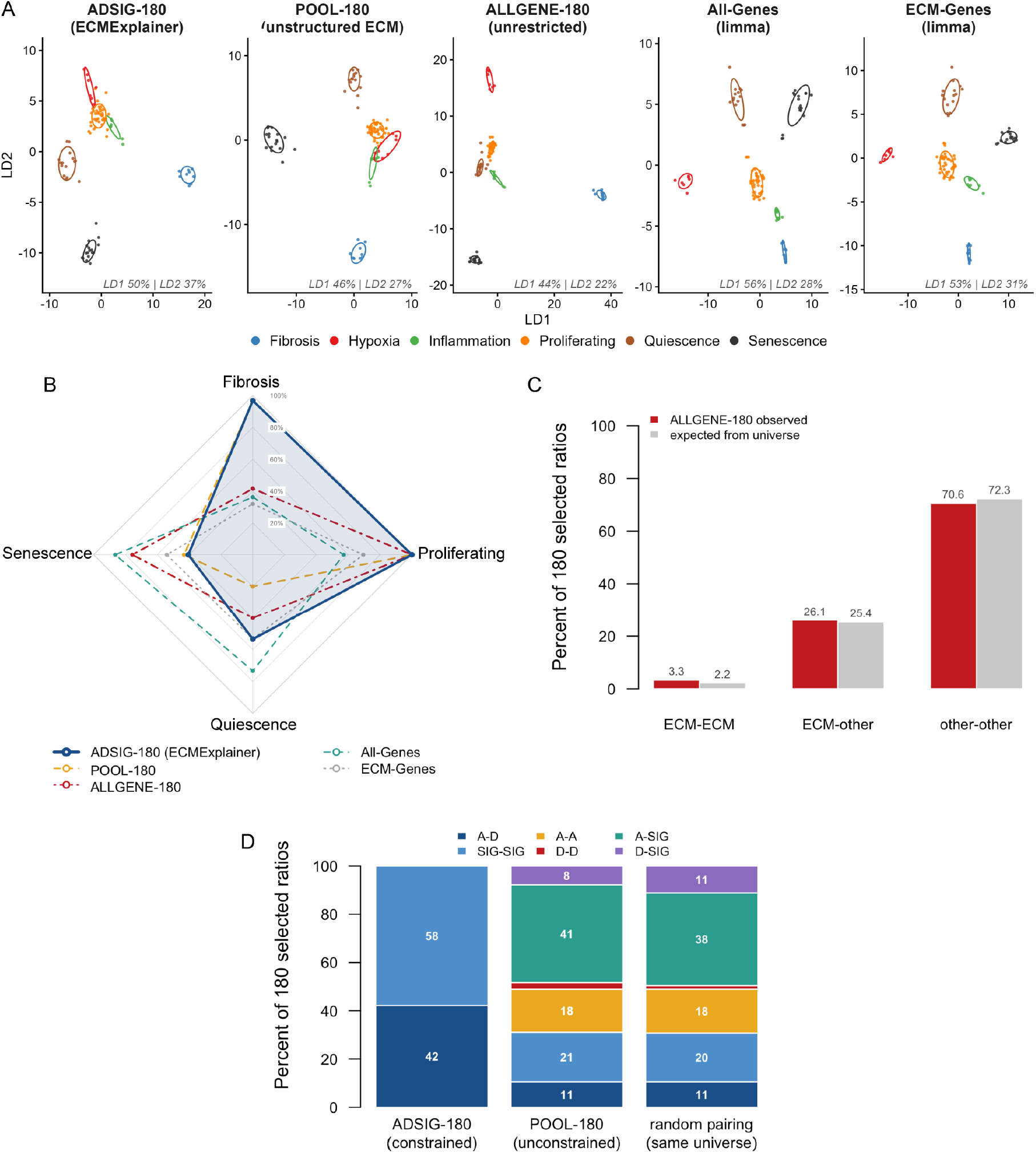
Benchmarking the ECMExplainer against competing classifiers. (A) LDA projections of the training samples for all five models (ADSIG-180 [ECMExplainer], POOL-180, ALLGENE-180, All-Genes, ECM-Genes), showing phenotype separation in LD1–LD2 space. (B) Radar chart of per-phenotype recall for each model across the 195 high-confidence scored validation samples (Fibrosis n=94, Proliferating n=49, Quiescence n=15, Senescence n=37). Each axis represents one phenotype: the outer ring = 100% recall. Balanced accuracy, reported in the main text, is the unweighted mean of the four axis values. ADSIG-180 (ECMExplainer, navy, solid line with shaded polygon) is the primary model; comparators are shown as dashed reference lines. Samples excluded from accuracy scoring (n=132: Supplementary Table 6) are retained for trajectory analysis only. (C) Composition of the 180 ALLGENE-180 ratios by whether each paired gene belongs to the curated ECM gene set (red, observed) against the proportion expected if 180 ratios were drawn at random from the same 2,000 gene top-variable universe (grey, expected). (D) Functional class of the two genes in each of the 180 selected ratios for the constrained ADSIG-180 panel (left), the unconstrained POOL-180 model (center) and the null-expectation for random pairing within the same 711-gene curated ECM universe (right).

To construct a high-confidence evaluation set, samples with ambiguous or experimentally undetermined phenotypes were excluded from the accuracy analysis and reported separately. Specifically, we excluded 132 samples in five categories: (i) 84 SRP102952 cardiac fibroblast baseline samples, whose ground-truth is undefined prior to TGF-β1 stimulation given the diseased nature of the human patients; (ii) 11 early senescence timepoints (≤4 days post-induction) where the senescent ECM program is not yet established; (iii) 3 samples with treatment status not clearly annotated in the source metadata; and (iv) 33 samples under pharmacological manipulation of unknown directional effect on the ECM (e.g. rapamycin, THZ1) or undergoing OSKM reprogramming, which are assessed by trajectory rather than hard phenotype label, and (v) one sample removed as a technical outlier before scoring. This yielded 195 scored samples across 18 datasets, with the 132 excluded samples retained for trajectory analysis only.

Across 18 independent cohorts (195 scored samples spanning four phenotypes), ADSIG-180 (the ECMExplainer) achieved the highest balanced accuracy at 72.7%, ahead of the size-matched POOL-180 (65.0%), ALLGENE-180 (64.3%), All-Genes (63.3%), and ECM-Genes (52.2%) (Figure 7B, Supplementary Table 6). This reflects the mean of the per-phenotype recalls, which weights each phenotype equally irrespective of how many samples it contributes. All three ratio panels excelled in their specificity, recalling 100% of Proliferating samples (49/49), while the two absolute-expression models performed substantially worse (All-Genes 57.1%, ECM-Genes 69.4%). Aggregate sensitivity across the three stress phenotypes (Fibrosis, Quiescence and Senescence; n=146) favored ADSIG-180 at 78.1%, ahead of POOL-180 (75.3%), All-Genes (52.7%), ALLGENE-180 (50.0%) and ECM-Genes (39.7%).

ADSIG-180 was therefore the only model to rank first on both stress sensitivity and balanced accuracy while retaining perfect control-state specificity. Its advantage over POOL-180 was concentrated in Quiescence, where ADSIG-180 recalled 8/15 samples (53.3%) against 3/15 (20.0%) for POOL-180. POOL-180 misassigned these samples to Proliferating, suggesting that unstructured ratio pools struggle to define the low-amplitude Quiescence state.

However, weighting each cohort equally rather than each sample reverses the ranking. ALLGENE-180 reached 88.0% mean per-dataset accuracy against 80.0% for ADSIG-180 and attained the highest accuracy in 14 of 18 datasets. This inversion is attributable to a single, large cohort. SRP102952 contributes 84 of the 94 Fibrosis samples and ALLGENE-180 recovered only 29 (34.5%) of them, assigning 55 to Proliferating. ADSIG-180 and POOL-180 both recovered 81/84 (96.4%). Because per-dataset gives this 84-samples cohort the same weight as a four-sample cohort, it conceals the largest failure in the benchmark. Outside SRP102952, every ratio model recalled all 10 remaining Fibrosis samples.

Second, Senescence was the limiting phenotype for both ECM ratio panels: ADSIG-180 recalled 15/37 (40.5%) and POOL-180 16/37 (43.2%), against 28/37 (75.7%) for ALLGENE-180 and 32/37 (86.5%) for All-Genes. Misalignments were predominantly to Proliferating (19/37 for ADSIG-180). ECM remodeling in senescence is heterogeneous and depends on the inducing stimulus and time since induction, so a fixed ECM ratio panel perhaps captures a narrower slice of senescent states than a transcriptome-wide signature ^68^. This is consistent with the senescent samples frequently occupying intermediate positions between the Proliferating and Senescence centroids in LDA space (as seen in Supplementary Figure 7–8). Further, the Senescence advantage of ALLGENE-180 was not derived from the matrix, as its ECM content arose by chance: 59 of 360 panel genes (16.4%) were ECM genes, indistinguishable from the 14.9% ECM content expected, and ECM genes were no more likely to be paired with one another, with only 6 ECM-ECM ratios observed (Figure 7C). Notably, no ECM-ECM ratio was selected for Senescence or Proliferating (Supplementary Table 6), instead, the Senescence quota was composed of immediate-early and MAPK-phosphatase genes (FOS, FOSB, DUSP4, DUSP6) together with mitotic and replication machinery (CENPF, CCNB2, SPC24, KIFC1) (Supplementary Table 6). ALLGENE-180 therefore reads stress amplitude and proliferative index rather than solely matrix program identity, indicating that these pathways provide a more universal route to detect senescence reliably than matrix composition does. This is therefore a boundary limitation of the ECMExplainer approach and motivates a senescence-specific extension of the ratio panel.

Since POOL-180 and ADSIG-180 achieved similar accuracy scores, we asked what functional class ratio pairs the unconstrained panel selected (Figure 7D). As the panels draw 180 ratios from the same 711 ECM genes, differing only in which functional class pairings can be made, the expected composition under random pairing can be obtained by enumerating all 252,405 possible gene pairs. POOL-180 matched the null, with just 56/180 ratios belonging to either A/D or SIG/SIG and sharing only 28 of its 180 ratios with ADSIG-180, without any significant loss in accuracy. Together, these comparisons show that ECM ratio-based representations provide the greatest advantage for Fibrosis, Proliferation and Quiescence, whereas Senescence relies more heavily on transcriptional programs beyond the matrix. These findings suggest that the ECM is an informative descriptor for some cellular states but not universally across all phenotypes, and the value of an ECM-restricted framework lies in interpretability and transferability rather than in uniformly superior discrimination.

## 4 Discussion

Here we establish a framework for describing ECM states based on coordinated matrisome gene expression. A key feature of this approach is the use of ratios between ECM gene modules, which capture relative changes in ECM composition while reducing technical and tissue-specific variation. This framework enables quantitative comparison of ECM remodeling across physiological and pathological conditions and provides a common language for characterizing ECM states.

The matrisome has been extensively categorized and annotated ^1–3^, and recent tools have extended that foundation to interaction inference (MatriCom, Lamba et al. 2025), spatial quantification (MatriSpace, Oshinjo et al. 2026), collagen architecture (Collagen Code, Zhang et al. 2023; CoDA, Nag et al. 202) and age-association or prediction (Matreotypes, Statzer et al. 2021; Plasma ECM clock, Coenen et al. 2026). These approaches are useful to inform us on what a matrisome profile contains or where it is located. However, the relationship between ECM profile and the phenotypic states of the cells that produce them remains poorly characterized. Instead of asking which ECM genes change under different cellular states, we asked whether, using functionally defined groups, the relative ordering of matrisome genes could place a given sample within a defined space of fibroblast ECM states. The ECMExplainer output is therefore not a label, but a coordinate, which allows intermediate and novel samples to remain interpretable.

The functional division of ECM genes into Assembly, Destruction and Signaling was imposed as a scaffold for pairing and interpretation, but the way the selected ratios behaved was itself informative. A ratio is only selected when the ordering of its genes differs between phenotypes, so what varies across all ratios is how the change arises. In 40% both genes moved in the same direction but by different amounts, tracking a shift in balance between two co-regulated genes, while a further 40% were dominated by a single partner and 20% were genuinely reciprocal. That reciprocal and concordant ratios together make up a large majority, suggests that phenotypes re-weight coordinated matrisome modules rather than switching one gene on and its partner off. Although the ECMExplainer only reads the direction of each comparison, the continuous log-ratio and the rank shift of each gene is retained, so any ratio of interest can be opened up at gene level. The transcriptional logic of these coordinated shifts remains unresolved. Perturbing individual pathways to ask which ratios move would turn a descriptive readout into a route to the transcriptional control of ECM state. These values will also aid future work to explore the transcriptional logic of these coordinated shifts.

Across 18 independent cohorts, matrisome expression was sufficient to assign unseen fibroblast transcriptomes to their correct reference state, with a balanced accuracy of 72.7%. Two comparisons show where this performance comes from. First, the gain lies in the encoding rather than the gene set. Using ECM genes read as absolute expression reached only 52.2% accuracy, so it is the rank-based representation, not the choice of genes, that transfers across cohorts. Second, the functional constraint we imposed did not add discriminative power. An unstructured ECM ratio panel drawn from the same ECM genes (POOL-180) shared only 28 of 180 ratios with ECMExplainer and showed no enrichment for A/D or SIG/SIG pairings, yet its balanced accuracy did not differ significantly. The A/D and SIG/SIG constraint therefore did not identify a privileged discriminative axis within the ECM, but added interpretability, where each of the 180 features reads as a specific shift in matrix assembly, turnover or signaling.

The ECMExplainer reached comparable per-dataset accuracy to a genome-wide ratio panel (ALLGENE-180, 88.0%; ECMExplainer, 80.0%) whose ECM content arose by chance, indicating that matrix-specific features are not required to reach this level of discrimination but also suggesting that the matrisome can capture information distinct from conventional transcriptomic analysis. These results establish the matrisome as a compact, interpretable and transfer-robust representation of fibroblast state. Whether it carries additional information that is genuinely additional to cell-intrinsic transcriptional programs is a separate question that our design was not built to answer, which we regard as an important question this work raises.

Interest in interpreting the ECM is growing for several reasons. Extracellular proteins are inherently more accessible than intracellular targets, and the extracellular space is more amenable to large therapeutic modalities such as antibodies, so the matrisome is attractive as both a diagnostic compartment and a drug target. Realizing this potential requires moving beyond collagens as surrogate markers to a broader set of matrisome-associated proteins and a wider range of phenotypes, which is what a multi-state ratio-based framework provides. In the context of restoring aged fibroblasts to more youthful, functional states, manipulation of the matrix may prove more tractable than manipulating cell-intrinsic programs, but this requires a readout that reports where in ECM space an intervention has moved a cell. Our pharmacological analyses illustrate this use case. Comparing two independent rapamycin-treated datasets identified FST/ITGA1 as a candidate pharmacodynamic ECM readout. Further, the TGF-β1 treated samples in GSE277437 that were also given the CDK7-inhibitor, THZ1, returned to a Proliferating ECMExplainer state from TGF-β1 alone which remained predicted as Fibrosis, even though the global transcriptome still placed TGF-β1 + THZ1 treated samples distinct from proliferating controls. Isolating the matrisome from genome-wide expression therefore does not merely reproduce the global response at lower dimension, it can report that the matrix program has been reversed while other programs have not. We therefore see the ECMExplainer as a candidate framework for screening compounds for the desired direction of ECM remodeling rather than for cytotoxicity or growth arrest alone.

Defining six homeostatic and stress states rather than a single contrast is what makes the ECMExplainer useful beyond the phenotypes it was trained on. Chronological aging and Hutchinson-Gilford progeria illustrate this point. Both are ‘aged’, yet they occupy different directions in ECM space, with progeroid fibroblasts aligning with fibrotic and inflammatory states, while chronological aging aligned with the quiescence-senescence axis. Neither a one-versus-one classifier nor a scalar age score can express that distinction, and it is the kind of statement a coordinate system naturally makes.

Senescence was the weakest phenotype for the ECMExplainer compared to the all-gene ratio model which we attribute to the heterogeneity in ECM remodeling depending on inducer and time point. That a rapamycin-responsive ratio was identified across multiple studies with different cell types and inducers within the Senescence ratio quota is striking and suggests that we were able to capture some universal matrisome features of Senescence. However, we admit a superior model would be a senescence-specific ECMExplainer, trained across inducers (replicative, oncogene-induced, genotoxic, oxidative and reprogramming-associated) and across time induction. This would allow that region of ECM space to be resolved into sub-states and would test directly whether an ECM-early and an ECM-mature senescent program are separable. Further, it would be worthwhile to generate ECMExplainer’s beyond human transcriptomes, such as mice, which would make it usable for intervention studies and genetic models, and cross-species comparisons to identify which functional balances are conserved properties of an ECM state.

ECMExplainer provides a quantitative and interpretable language for ECM state that transfers across datasets without retraining. Broadening the reference space to further states, inducers and species and identifying the regulators that set these ratios remain necessary next steps and will bring us closer to the aim of a complete functional interpretation of ECM gene expression. What the framework delivers now is matrisome remodeling made measurable and comparable between conditions, interventions and forms of aging.

## 5 Limitations

The training atlas is small. It contains 96 samples from 10 studies, and two of the six phenotypes (Hypoxia and Inflammation) are each represented by only two studies. This explains the gap between external validation (balanced accuracy 72.7%) and leave-one-study-out cross validation on the training atlas (47.9%). We interpret this gap as data scarcity rather than overfitting, because the model generalized well to 18 cohorts it had never seen, but the interpretation cannot be confirmed until the atlas is larger. For the same reason, accuracy did not differ significantly between the 180-ratio panel and any other size from 60 to 247 ratios, so cross-validation on this atlas provides no basis for tuning panel size. Expanding to at least three independent studies per phenotype is the first priority for a second version.

The phenotype space is incomplete. Hypoxia and Inflammation were not validated on external data, because no suitable public fibroblast cohort was available at submission, so their behavior on independent samples remains untested. Other stress states, such as apoptosis, are absent altogether because suitable datasets were not available. Predictions for conditions outside the six reference states should therefore be read as positions in the ECM space rather than as class labels.

All data are from fibroblast monolayers *in vitro*. Tissue-level or single-cell data contain non-fibroblast cells whose matrisome expression differs, and ratio-based predictions have not been tested in that setting.

Finally, the ECMExplainer reads transcripts, not matrix. It does not capture ECM protein abundance, post-translational modification, cross-linking or spatial organization, which are properties that define matrix function. Integrating the transcriptional states defined here with proteomic and imaging measurements of the same samples is the direct next step and will show how far transcriptional ECM state reflects the matrix itself.

## 6 Methods

### 6.1 Data Acquisition & preprocessing

To build the ECMExplainer, ninety-six RNA-sequencing samples were aggregated from ten independent human fibroblast studies deposited in the NCBI Gene Expression Omnibus (GEO): GSE63577 ^69,70^, GSE74324 ^71^, GSE75643 ^72^, GSE102674 ^73^, GSE117337 and GSE117444 ^54,74^, GSE139963 ^75^, GSE221693 ^76^, GSE281533 ^77^, and GSE295622 ^78^. Transcripts were quantified with Salmon v1.10.2^79^ against a transcriptome index built from the Ensembl release 111 Homo sapiens GRCh38 cDNA FASTA (Homo_sapiens.GRCh38.cdna.all.fa.gz) using salmon index with default parameters. Paired-end reads were quantified with salmon quant in mapping-based mode using flags --seqBias --gcBias --validateMappings; transcript-level estimates were summarized to gene level with tximport v1.30.0 ^80^ using a GENCODE v46 tx2gene map and countsFromAbundance = "lengthScaledTPM".

The resulting gene-by-sample count matrix for all samples was integrated with curated sample metadata for downstream modeling of ECM-associated transcriptional responses. The combined dataset included 96 samples across six distinct cellular phenotypes: Proliferating, Senescence, Quiescence, Fibrosis, Inflammation, and Hypoxia. To cover a comprehensive view of tissue aging and extracellular matrix remodeling, the datasets included diverse cell lines including human neonatal foreskin fibroblasts (BJ and HFF), human fetal lung fibroblasts (IMR90 and WI38) and iPSC-derived cardiac fibroblasts.

For quality control, Principal Component Analysis (PCA) was plotted for each individual dataset using all available genes to confirm phenotypic separation and to confirm there are no outlier samples that deviated from the main cluster. Each dataset included Proliferating samples. After gene symbol harmonization across platforms and intersection of measured genes, a common feature space of 19,861 genes was retained across all 96 samples.

In addition, PC QC (top 2,000 variable genes and ECM-subset) was performed pre-and post-batch-correction using limma::removeBatchEffect with study as the batch factor and phenotype as the protected design (∼phenotype + batch), confirming phenotype separation after correction.

To confirm that all 96 training samples represented a common fibroblast lineage despite their diverse phenotypic states, per-sample lineage scores were computed as the mean log_2_(CPM + 1) expression of curated marker panels on the post-batch-correction matrix. The fibroblast panel comprised COL1A1, COL1A2, COL3A1, PDGFRA, PDGFRB, VIM, DCN, LUM, FAP, S100A4, THY1 and FN1, drawn from cross-tissue single-cell fibroblast atlases ^21,22^. Non-fibroblast lineage panels were defined from the Tabula Sapiens human cell atlas (Tabula Sapiens Consortium et al. 2022) as epithelial (EPCAM, KRT8, KRT18, KRT5, CDH1), endothelial (PECAM1, CDH5, VWF, KDR) and immune (PTPRC, CD3D, CD68, CD14, ITGAM). Smooth-muscle markers were restricted to the contractile-phenotype set defined by Owens et al. (2024) (MYH11, CNN1, TAGLN), and pluripotency was assessed using the canonical reprogramming factors (POU5F1, NANOG, SOX2; Takahashi et al. 2007). A sample passed the fibroblast-dominance gate if its fibroblast-panel score exceeded the highest non-fibroblast-panel score. All 96 samples passed, with a minimum margin of 0.54 log_2_ CPM units.

### 6.2 ECMExplainer curated Gene list

ECMExplainer gene panel construction. The ECMExplainer panel was assembled as the union of the Human Matrisome Portal masterlist (Naba et al., 2012; 1,027 genes) and nine ECM-focused pathway gene sets retrieved from MSigDB v2026.1.Hs (Liberzon et al., 2011, 2015), covering collagen biosynthesis and fibrillogenesis (REACTOME_COLLAGEN_FORMATION, GOBP_COLLAGEN_FIBRIL_ORGANIZATION), basement-membrane organization (GOBP_BASEMENT_MEMBRANE_ORGANIZATION), glycosaminoglycan and hyaluronan metabolism and turnover (KEGG_GLYCOSAMINOGLYCAN_BIOSYNTHESIS_HEPARAN_SULFATE, KEGG_GLYCOSAMINOGLYCAN_BIOSYNTHESIS_CHONDROITIN_SULFATE, KEGG_GLYCOSAMINOGLYCAN_DEGRADATION, REACTOME_HYALURONAN_METABOLISM), and cell–matrix signaling (KEGG_ECM_RECEPTOR_INTERACTION, GOBP_COLLAGEN_ACTIVATED_SIGNALING_PATHWAY). A gene was included if it was a member of at least one source set; no gene was added on the basis of literature alone. Symbols were harmonized to HGNC nomenclature, with withdrawn or ambiguous aliases resolved manually, giving a union of 1,175 unique genes.

Thirty-six genes were then excluded under a uniform ECM-specificity criterion, applied independently by two curators and reconciled by discussion: a gene was removed if it appeared in its source sets only as an intracellular effector, transcription factor, membrane transporter, cytoskeletal adaptor or oxygen sensor whose characterized substrates are not secreted matrix, or as a clear pathway-annotation artefact (the synaptic-vesicle glycoproteins SV2A/B/C). Seven of these derived from the matrisome masterlist itself (EGLN1–3, P4HTM, OGFOD1, OGFOD2, NGLY1); all are matrisome-associated ECM Regulators annotated by 2-oxoglutarate-dependent dioxygenase or glycanase domain homology, but with characterized intracellular substrates (HIF-α, ribosomal protein RPS23, ERAD glycoproteins). No core matrisome gene was excluded, and the collagen-modifying enzymes P4HA1–3, P4HB and PLOD1–3 were retained. Per-gene exclusion reasons are given in Supplementary Table 1 (sheet "Excluded_Genes").

Each of the 1,139 retained genes was assigned to a single functional module, Assembly (n = 412), Destruction (n = 119) or Signaling (n = 608), reflecting its dominant contribution to matrix architecture as described by its existing database annotations. Assignments were derived programmatically from the annotations already attached to each gene: its Matrisome Portal division and category where available, and otherwise its membership of the contributing MSigDB pathway sets. Because these source annotations do not map one-to-one onto the three processes, the category "ECM Regulators", for example, contains proteases, endogenous protease inhibitors and collagen-modifying enzymes, which act in Assembly and Destruction respectively, the programmatic assignments were then manually inspected, and corrected where the inherited label conflicted with the gene’s described function. Ten genes were reassigned on this basis, principally the procollagen N-and C-proteinases and their enhancers (ADAMTS2, ADAMTS3, ADAMTS14, BMP1, TLL1, TLL2, PCOLCE, PCOLCE2), which are proteolytic by family but act in fibril assembly, and the hyaluronidases CEMIP and SPAM1. Review was targeted at families where annotation and function were expected to diverge rather than applied exhaustively to all 1,139 genes. Per-gene module, matrisome division and category, contributing source sets and curation notes are given in Supplementary Table 1.

Module assignment therefore reflects the functional descriptions currently attached to each gene in its source annotations, together with the deliberate simplification of assigning one gene to one module. Many matrisome proteins are multifunctional, and some assignments would differ if a different annotated activity were taken as primary. The partition is intended as a pragmatic functional scaffold for constructing gene-pair ratios, not as a definitive functional ontology; it is provided in full so that alternative assignments can be applied by others.

The ECMExplainer panel was intersected with genes expressed in the non-batch-corrected training matrix (CPM > 1 in at least as many samples as the smallest phenotype class [n = 7, Hypoxia], and variance > 0.005 across the raw logCPM matrix), retaining 711 ECM genes (Assembly n = 303, Destruction n = 88, Signaling n = 320) as the active modelling universe from which candidate gene-pair ratios were drawn. The corresponding filter applied to the limma batch-corrected matrix, used for the descriptive ECM-space analyses in Figures 1B–D, retained 895 of the 923 ECMExplainer genes detected in all 96 samples (28 genes, 3.0%, discarded). The two gene universes therefore differ by design: 895 genes support batch-corrected magnitude-based visualization, while the 711-gene uncorrected universe supplies candidate gene pairs for ratio selection.

To characterize the macroscopic ECM transcriptional state of each training phenotype, we computed per-sample Assembly and Destruction module scores as the mean batch-corrected log2 CPM across the retained genes of each module, and the Net ECM Score as their difference, S_net = S_A − S_D, as a summary of matrix-deposition balance. Differences in Net ECM Score across phenotypes were visualized as median-ordered violin plots, and the underlying Assembly and Destruction components were displayed as a bivariate scatter with per-phenotype 85% confidence ellipses.

### 6.3 k-TSP Ratio Selection

#### Sign-encoded ratio features

ECMExplainer is built on sign-encoded gene-pair ratios following the k-Top Scoring Pairs (k-TSP) paradigm ^28,29^. For each gene pair (A, B) and each sample *i*, the feature is the within-sample rank order of the two genes: x_{AB,i} = sign(logCPM_{A,i} − logCPM_{B,i}) ∈ {−1, 0, +1}. Any per-sample additive offset applied identically to both genes (library-size rescaling, TMM) cancels inside the difference, and any monotone within-sample transform applied to all genes (variance-stabilising transformation -VST) preserves their rank order, so x_{AB,i} is invariant to these confounders by construction. Batch effects, however, generally include a gene-specific component that does not cancel; rather than model it explicitly, we rely on cross-validated pair selection (leave-one-study-out, below) to down-weight pairs whose ordering is unstable across studies, in deliberate contrast to the magnitude-based benchmarks.

#### Pair families and panels

Ratios are constructed within two functional modules: A/D (Assembly/Destruction, asymmetric cross-module pairs) and SIG (Signaling/Signaling, symmetric pairs with alphabetical ordering). Two panels were built: an unstructured control (POOL) drawing from all expressed, variance-filtered ECM genes, and the primary structured model (ADSIG) drawing from the AD and SIG families only. A third ratio panel, ALLGENE, was built that serves as another unrestricted control, drawing from the 2,000 most variably expressed genes.

#### Two-stage candidate scoring and panel assembly

Within each panel, candidate ratios were scored in two stages. First, discriminative variance was quantified as η² for the continuous log-ratio defined as log_2_(CPM_a_ + 1) − log_2_(CPM_b_ + 1), which reflects the proportion of its variance explained by the phenotype. Ratios with η² ≤ 0.10 were discarded. Surviving ratios were then scored by TSP delta, defined as the largest one-versus-rest difference in the frequency with which the numerator ranked above the denominator across the six phenotypes. Each ratio was assigned to the phenotype yielding its maximum TSP delta. Thus, η² served as a variance-based admission filter, while TSP delta supplied a phenotypic-specific ranking statistic for panel selection. Ratios were considered in descending TSP delta order. A ratio was added if its tagged phenotype had not yet reached its quota of N/6 and neither constituent gene had already been selected. Each gene was therefore used in at most one ratio and genes were allocated to the most discriminative ratios first.

#### Classifier construction

The 180 × 96 sign matrix was reduced by a single joint PCA (centered and scaled). Components explaining ≥95% of cumulative variance were retained, up to a maximum of ten; where ten components did not reach 95%, the cap was applied. Because features are sign-encoded {−1, 0, +1}, scaling normalizes each ratio to unit variance prior to rotation. The retained PC scores were classified by six-class LDA (MASS::lda), producing a five-dimensional discriminant space with one training centroid per phenotype.

#### Panel size and cross-validation

Panel size was assessed from 60 ratios upward in increments of 30 ratios, corresponding to five additional ratios per phenotype at each step. The A/D arm was bounded by the 88 expressed Destruction genes and the SIG arm by 160 Signaling gene pairs, giving a theoretical maximum of 248 ratios, of which 247 were achievable after η² filtering and phenotype-quota constraints. Two cross-validation protocols were used and are reported separately. Leave-one-out cross-validation (MASS::lda, CV = TRUE) was applied to the LDA classifier on the fixed final panel and PCA rotation,

both of which were estimated once on all 96 samples; the LOOCV figure therefore reflects classifier stability on a pre-selected feature representation. LOSO and external validation are reported as informative metrics. Nested leave-one-study-out cross-validation (LOSO) was used as the generalization estimate: for each held-out study, the full pipeline; variance filter, η² candidate pool, ranking, greedy selection, PCA rotation and LDA, was re-estimated on the remaining nine studies, with no features, rotation or scaling shared across folds.

#### LOSO performance

ADSIG LOSO performance did not differ significantly with panel size across the sweep (43.8–52.1%). The final version ADSIG-180 (ECMExplainer) attained 47.9% nested LOSO accuracy. Since overall accuracy is dominated by the larger classes, we also report per-phenotype recall: Proliferating 85.0%, Quiescence 46.7%, Fibrosis 22.2%, Senescence 17.6% Hypoxia 0% and Inflammation 0%, giving a balanced accuracy of 28.6%. This is driven by Hypoxia and Inflammation, the two phenotypes contributed by only two source studies each. Withholding either study leaves a single remaining study to define that centroid, and per-fold recall for both was zero. Restricting to the four phenotypes represented by ≥3 source studies (Fibrosis, Proliferating, Quiescence, Senescence) gives 56.8% accuracy and 42.9% balanced accuracy (Supplementary Table 3). Panel sizes were compared using paired exact tests on discordant classifications from the same LOSO samples (McNemar’s test). No panel size different significantly from the ADSIG-180. We therefore retained the 180-ratio panel because it provides 30 ratios per phenotype increasing redundancy against genes that may be absent in external cohorts. The final ADSIG-180 panel comprises 76 A/D and 104 SIG ratios.

#### Permutation test of individual ratios

To test whether the individual selected ratios separated their tagged phenotype beyond chance, phenotype labels were randomly permuted 2,000 times per ratio and the maximum one-versus-rest sign-frequency separation was recalculated. Empirical p-values were adjusted across all 180 ratios using the Benjamini-Hochberg procedure; ratios with FDR < 0.05 were classified as discriminative.

#### Behavior of ratio-constituent genes

Gene-level expression behavior was quantified using within-sample percentile ranks rather than absolute expression values. For each sample, genes were ranked across all expressed genes and expressed as percentile ranks. This transform is insensitive to library size, normalization method and sample-level technical shifts. Proliferating samples, which are present in all ten training studies, served as the within-study reference. For each of the five non-Proliferating phenotypes, the contrast was that phenotype versus Proliferating samples from the same study. For the 30 ratios selecting the Proliferating phenotype, the contrast was reversed to Proliferating versus all non-proliferating samples from the same study. For each gene we computed the difference in mean percentile rank between the two groups within each contributing study, then combined by inverse-variance fixed-effect meta-analysis, reporting the pooled shift with its 95% confidence interval. The same procedure was applied to the ratio itself, defined per sample as rank(numerator) – rank(denominator).

Each ratio was then assigned to one of four behavior classes in two steps. First, if both genes shifted in the same direction, the ratio was classes as a concordant shift, since it then reports a change in the balance between two co-regulated genes. Second, for ratios whose genes shifted in opposite directions, we computed the numerator’s share of the total movement, *c_A_ = |Δ_A_|/*(|*Δ_A_*| *+*|*Δ_B_*|), where *Δ_A_* and *Δ_B_* are the pooled shifts of the numerator and denominator. When *c_A_* ≥ 0.70, the ratio was classed as Gene A dominant. For ratios with *c_A_* ≤ 0.30), the ratio was Gene B dominant. For remaining ratios, 0.30 < *c_A_* < 0.70, the ratio was classified as reciprocal. The one-versus-rest (OvR) score is the frequency with which the numerator ranks above the denominator in the selecting phenotype minus the frequency in all other samples (-1 to +1). All per-ratio, per-gene and per-study estimates are given in Supplementary Table 3.

#### Magnitude baselines and deployment

Two magnitude-based controls were trained on the same 96-sample matrix, differing from the ratio panels only in feature encoding and universe: an All-Genes model (18,957 genes with logCPM variance > 0.005) and an ECM-Genes model (the 895 ECMExplainer genes meeting the same variance criterion; note this is a slightly larger universe than the 711-gene ratio candidate pool, which additionally required CPM > 1 in ≥ 7 samples, so the baseline is not disadvantaged by gene coverage). Both were batch-corrected with limma::removeBatchEffect (study as batch, phenotype-protected design: ∼phenotype + batch) prior to PCA → LDA, as absolute logCPM is not batch-invariant.

Ratio panels and gene baselines were deployed under different test-time contracts. Ratio panels required no test-time normalization: each sign feature x_{AB,i} = sign(logCPM_{A,i} − logCPM_{B,i}) is computed within a single sample and projected through the stored training PCA (predict.prcomp) and stored LDA; ratios with a missing gene were set to sign 0. Gene baselines were projected by reference-projection. For each gene g and validation sample i, the within-cohort z-score (x_{g,i} − μ^val_g)/σ^val_g was computed using only that cohort’s samples, then divided by the stored training scale σ^train_g and projected through the stored PCA loadings by direct matrix multiplication; predict.prcomp was not used, to avoid re-applying the training center and scale over the within-cohort standardization. Genes with zero within-cohort variance, and cohorts too small to estimate a within-cohort SD, were assigned a scale of 1 (centered only); genes absent from a cohort were set to the neutral standardized value 0.

Reference-projection used only the target cohort’s own samples and no training or cross-cohort data. It is transductive: μ^val and σ^val are cohort-level statistics, so a sample cannot be scored in isolation. This procedure raised cross-study LOSO accuracy (All-Genes 28→45%; ECM-Genes 40→49%) while leaving LOOCV and model ranking unchanged. Because the frozen rotation was fitted on training-scaled inputs, cohort-level rescaling aligns the projection axes only when per-gene dispersion is comparable across cohorts.

### 6.4 Validation Projection Pipeline

ECMExplainer was applied to 18 independent public validation cohorts spanning senescence, quiescence and fibrosis: SRP113329, SRP117883, SRP104418, SRP121031, SRP034541, SRP040745, SRP098713, SRP136071, SRP017142, ERP021140, SRP153724, SRP153205, SRP052706, SRP045867, SRP102952, GSE277437, GSE302114 and GSE273882. Two further cohorts were analyzed exploratorily rather than for accuracy assessment: GSE113957 (chronological and progeroid aging), from which the ECM Aging Score was derived, and GSE165176 (partial OSKM reprogramming), to which that score was applied without refitting. Both are assessed by centroid-distance trajectory rather than hard phenotype label and contribute to no reported accuracy figure.

Raw counts or author-normalized matrices were loaded and converted, where required, to log₂(CPM + 1). Gene identifiers were mapped to HGNC symbols with AnnotationDbi/org.Hs.eg.db; where a symbol mapped to multiple rows, the row with the highest variance in that cohort’s log₂-CPM matrix was retained, so the surviving probe for a given symbol is cohort-specific. Because the ratio encoding is sign(log₂X_A – log₂X_B), any strictly per-sample scaling cancels within the sign; log₂-CPM and log₂-RPM inputs are therefore exchangeable, differing from true log₂-CPM only by a per-sample additive constant. Length-normalized units (RPKM, FPKM, TPM) are not exchangeable with CPM, because the per-gene length factor shifts the two ratio members unequally and does not cancel. No validation cohort scored for accuracy used such units. One exploratory cohort, GSE113957, is deposited on GEO only as FPKM and was analyzed as log_2_(FPKM+1). As the length factor is constant across samples for a given ratio, within-cohort age slopes are unaffected and centroid distances keep their direction. Cohorts deposited as log₂-RPM (GSE165176) were confirmed to be per-sample-scaled RPM and projected without further renormalization.

The scorable unit is the ratio pair, not the individual gene. A ratio (A, B) was computed only when both members were present in the cohort; the continuous log-ratio r = log₂X_{A} − log₂X_{B} was formed and encoded as x = sign(r) ∈ {−1, 0, +1}. Ratios in which either member was absent were neutralized to 0, contributing neither a positive nor a negative vote — the deployment analogue of the training missing-ratio rule. All cohorts exceeded 85% pair coverage (the fraction of the 180 ratios with both genes present), 5 at 100%, so no validation metric is materially influenced by neutralized ratios.

Each sample’s sign vector was projected through the frozen training PCA: the stored training per-gene center and scale were applied and the vector rotated by the stored training eigenvectors (predict.prcomp on the saved prcomp object), retaining the same fixed number of components used at training (≤10), then classified by the stored LDA. The rotation columns and component count are the stored training values and are not re-estimated on validation data. Reuse of the training-estimated center and scale is exact here, unlike the cohort-centered gene baselines, because the sign features are per-sample batch-invariant and carry no cohort-specific location, so no test-time recentering is required. Consequently, all validation samples occupy the training coordinate system, and distances, classifications, and posterior probabilities remain directly comparable.

#### LDA classification and posterior probabilities

The validation PC matrix was passed to the trained LDA model. For each sample this produced a posterior probability across the six phenotypes (Proliferating, Quiescence, Senescence, Fibrosis, Inflammation, and Hypoxia), a hard-assigned label corresponding to the highest posterior, and a set of discriminant coordinates. Posterior probabilities were plotted as per-sample stacked bars, grouped by experimental condition.

### 6.5 Centroid geometry, distance heatmaps and trajectory angles

#### Centroids and distances

Training phenotype centroids μ_p were computed as the per-class means of the training LDA scores from the frozen model, in the sphered LDA discriminant space in which within-class covariance is approximately isotropic; plain Euclidean distance in this space is therefore equivalent to the Mahalanobis distance the classifier uses, which justifies its use for both distance and centroid geometry. All six μ_p are the fixed training centroids obtained from the stored PCA→LDA scores without refitting, and every validation sample is placed against them in the same frozen coordinate system, so distances are directly comparable across cohorts.

#### Distance heatmaps

For every validation sample we computed the Euclidean distance to each of the six training centroids. Per-condition median distances were rendered as a six-column distance heatmap. Because a median distance measures proximity only and discards direction, the heatmap is interpreted strictly as a proximity summary; directional (trajectory orientation) claims are carried separately by the trajectory panels.

#### Trajectory geometry — dose and time series

For intervention (vehicle → treated dose series) and reprogramming (Baseline → iPSC over time) comparisons, each condition was represented by its medoid — the real sample minimizing total distance to the other samples in that condition — and arrows connect successive medoids in the two-centroid plane spanned by the Proliferating and Senescence training centroids. Medoids (actual sample coordinates rather than means of few points) were used to avoid the instability of small per-condition averages in discriminant space; the number of samples underlying each condition is reported per arm, and a triangle-inequality check (distance-to-Senescence + distance-to-Proliferating ≥ the Senescence–Proliferating centroid separation) confirms every plotted point respects the LDA geometry. For the reprogramming time course the deployable readout is the median LDA distance to the Proliferating, Quiescence and Senescence centroids as a function of day.

#### Trajectory orientation — chronological aging protractors

For the chronological-aging cohort, the observed aging direction was defined entirely within validation as v = μ_Aged^val − μ_Young^val (or μ_HGPS^val − μ_Young^val for progeroid aging), with the Young validation centroid μ_Young^val serving as the common origin. For each training phenotype p, the candidate direction μ_p − μ_Young^val was constructed from that same origin, and the angle between v and each μ_p − μ_Young^val was computed as θ = arccos( (v·d)/(ǁvǁǁdǁ) ), reported in degrees; because both vectors share the origin μ_Young^val, the angle measures purely whether the empirical aging trajectory points toward a given phenotype, independent of where the cohort baseline sits. Arrow length was scaled to phenotype distance and rendered as protractor figures. To quantify stability given small per-group sample sizes, angles were bootstrapped (2000 resamples, resampling within the Young and Aged/HGPS groups), and 95% confidence "fans" were drawn from the 2.5th–97.5th percentiles; the protractor is generated only when each group has ≥2 samples.

#### Trajectory protractor analysis and bootstrap confidence intervals

For each age contrast (Young → Aged and Young →HGPS), the reference aging trajectory was defined in LDA space as the vector from the Young-group centroid to the Aged-or HGPS-group centroid. For each training phenotype, a phenotype vector was drawn from the Young centroid to that phenotype’s training centroid, and its angular alignment with the reference trajectory was computed as the arccosine of the cosine similarity between the two vectors (0° = identical direction, 90° = orthogonal, 180° = opposite). To estimate uncertainty, we applied a nonparametric bootstrap (n = 2,000 iterations; fixed seed for reproducibility): at each iteration the Young group and the target group (Aged or HGPS) were independently resampled with replacement, their centroids recomputed, and the reference trajectory and all phenotype angles re-derived from the resampled centroids. The reported angle is the median across bootstrap replicates, and the 95% confidence interval is the 2.5th–97.5th percentile of the bootstrap distribution (percentile method). Arrow lengths in Figure 5D are the Euclidean distances from the Young centroid to each phenotype centroid, normalized to the maximum such distance. All computations were performed in R using base linear-algebra routines and the MASS package for the LDA projection.

### 6.6 Senescence kinetics and pharmacological interventions

#### Trajectory arrows

Each validation sample was assigned two scalar coordinates in LDA space: its Euclidean distance to the frozen training Senescence centroid (D_Sen) and to the frozen training Proliferating centroid (D_Prolif). Each induced-senescence condition was represented by its medoid, the sample minimizing the summed Euclidean distance to all other samples of that condition, computed in the full multidimensional LDA space rather than in the two-dimensional distance projection. Using the medoid guarantees that every plotted point corresponds to a real sample and therefore satisfies the triangle inequality D_Sen + D_Prolif ≥ ǁμ_Sen − μ_Prolifǁ, which independent per-axis medians can violate. Arrow segments connect successive medoids to trace the rapamycin dose-response (untreated → 1 nM → 10 nM rapamycin) from the induced-senescence baseline toward the Proliferating anchor, drawn solid for the RAS-induced arm and dashed for the OSKM-induced arm. Gold-triangle at (0, d) and (d, 0), where d = ǁμ_Sen − μ_Prolifǁ is the separation between the two training centroids, mark the positions a sample would occupy if it coincided exactly with the Senescence or Proliferating centroid. Because this two-anchor plane collapses the five-dimensional discriminant space and cannot resolve movement orthogonal to the Senescence–Proliferating axis, the arrows are presented as a schematic. The reversal is quantified independently by the signed scalar D_Prolif − D_Sen (positive values indicate greater proximity to Senescence), reported per condition alongside per-arm sample size.

#### Rapamycin ratio concordance scatter

For every ratio j and each rapamycin dose d ∈ {1 nM, 10 nM}, we computed the mean shift in the continuous log₂ ratio relative to the untreated arm: Δ_RIS = r̄(RAS+d) − r̄(RAS) and Δ_OSKM = r̄(OSKM+d) − r̄(OSKM), and plotted Δ_OSKM against Δ_RIS per dose. Each ratio was assigned to one of four mutually exclusive classes. Ratios with min(|Δ_RIS|, |Δ_OSKM|) ≤ 0.05 log₂ units (equivalent to a ≈3.5% change in the underlying ratio) were classed as sub-threshold and excluded from concordance scoring. Above-threshold ratios were classed as concordant "Up in both" or "Down in both" when sign(Δ_RIS) = sign(Δ_OSKM), or "Discordant" when the signs disagreed. Percent-concordant was computed over scored (above-threshold) ratios only, so sub-threshold ratios were excluded from both the concordant and discordant counts rather than defaulting to discordant. Of the 180 panel ratios, 164 were scored at 1 nM and 163 at 10 nM; the remainder were sub-threshold or had an absent constituent gene in this dataset. We report Spearman ρ as the primary concordance statistic, as it is robust to the leverage of the few large-magnitude ratios that are also labelled; Pearson r is reported both across all ratios and across the sub-threshold-excluded set to expose any tail dependence. The 0.05 floor is an absolute cutoff on the log₂ scale; because per-arm sample sizes are small (n reported per dose in the figure), it should be interpreted as a descriptive effect-size filter rather than a noise-calibrated significance threshold. The top 6 concordant and top 4 discordant ratios per dose, ranked by |Δ_RIS| + |Δ_OSKM|, were labelled.

#### Kinetic panels

Each human cardiac fibroblast condition was assigned a label-level population-doubling (PD) value from the source study (EarlyPass = 30, LatePass = 52, RapaCtrl = 45, OldRapaMR = 85; Azar et al. 2018); these are literature values, not per-sample measurements. Rapamycin in this cohort was applied chronically at 1 nM throughout culture, and methionine restriction at 1 mg/L L-methionine (97% restriction), following the source protocol. Two trajectories were defined on a shared PD30 (EarlyPass) anchor: Natural Aging (EarlyPass → LatePass) and chronic Rapamycin (EarlyPass → RapaCtrl → OldRapaMR). MetR and RapaMetR (both PD45) were omitted from the kinetic axis to avoid overplotting at a shared x-coordinate; the fixed-PD drug contrast is shown in the centroid map and distance heatmap (Figure 3B). For each condition we computed the mean Euclidean distance, in the sphered LDA discriminant space, from projected validation coordinates to each of the frozen Proliferating, Senescence, Fibrosis, and Inflammation training centroids, plotting distance against PD with mean ± SE and n per knot. Because slopes depend on imputed PD coordinates, quantitative rate comparison is restricted to the overlapping PD 30–52 window (shaded); OldRapaMR (PD85) anchors an extrapolated segment no natural-aging point can validate. "Drift" denotes movement in classifier space, not literal ECM-composition change.

#### Cross-study rapamycin signature

To ask whether the rapamycin-responsive matrisome axis is shared across mechanistically distinct senescence programs, we compared rapamycin-associated ratio changes in Ras-induced (SRP117883) and replicative (SRP104418) senescence. For each of the 180 ADSIG-180 k-TSP ratios we computed, independently within each study, a treatment contrast: Δ_RIS = mean(RAS + 10 nM rapamycin) − mean(RAS) in SRP117883, and Δ_Rapa = mean(RapaCtrl) − mean(LatePass) in SRP104418. The 10 nM dose was specified a priori. Each Δ is a within-study difference, so any gene-specific offset that is constant within a study cancels; because the two Δ’s originate in different cohorts, however, their agreement is an explicitly cross-study quantity not protected by the within-arm cancellation that safeguards the single-study analyses, and we interpret it accordingly. We note the two contrasts are not identical operations (a within-study dose shift versus a treated-minus-late-passage difference); we therefore report shared ratio-level directionality rather than an identical rapamycin displacement. Ratios were merged on shared IDs (inner join; all 180 panel ratios were computable in both cohorts). Concordance was scored on ratios exceeding an absolute floor of |Δ| = 0.05 log₂ in both axes (sub-threshold ratios excluded from both numerator and denominator) and summarized by Spearman ρ (primary) and Pearson r (secondary). As a drug-specificity control, we repeated the comparison using Δ_MetR = mean(MetR) − mean(LatePass) from the same replicative cohort; greater rapamycin–rapamycin than methionine-restriction–rapamycin concordance indicates a rapamycin-specific rather than generic cross-study axis. Labelled ratios are the top 5 concordant and top 3 discordant by |Δ_Replicative| + |Δ_RIS|.

### 6.7 TGF-β1 Fibrotic Response

#### Paired patient trajectories

Patient-matched Baseline and TGF-β1-treated primary cardiac fibroblasts (SRP102952; n = 84 patients, 168 samples) were projected into ECMExplainer LDA space, and per-patient distances to the Fibrosis and Proliferating centroids were computed from a single RNA-seq sample per patient per condition, so reported values reflect individual measurements rather than averages across replicates. Baseline and TGF-β1-treated samples were normalized together within a single log₂-CPM transform prior to PCA/LDA projection, preserving the within-patient contrast, and baseline-to-treated arrows were then drawn per patient with gold-triangle anchors marking the frozen training Fibrosis and Proliferating centroids, preserving pairing that a population-level scatter would otherwise collapse. To formally test this shift, we treated each patient as their own control and applied a paired Wilcoxon signed-rank test to the per-patient change in Fibrosis-centroid distance following treatment (ΔD_Fibrosis = D_Fibrosis,treated − D_Fibrosis,baseline). Because the two-dimensional (D_Fibrosis, D_Proliferating) plane used for visualization can distort true displacement in LDA space, samples on opposite sides of the Fibrosis–Proliferating axis can appear as similar-length arrows despite moving in opposite true directions, we also computed the paired test on the full six-centroid distance profile, which confirmed the same directional result and ruled out an artifact of the two-anchor projection.

#### Waterfall of per-patient Euclidean shift

For each patient we computed ΔD_Fibro = D_Fibrosis(TGFβ) − D_Fibrosis(Baseline) — negative values indicate a move toward the Fibrosis centroid — and displayed these shifts as a rank-ordered waterfall bar chart colored by direction (Closer-to-Fibrosis in red, Further-from-Fibrosis in blue), annotated with the percentage of patients moving fibrotic and the paired two-sided Wilcoxon p-value. This visualization reveals both the average effect and the degree of patient-level heterogeneity in TGF-β1 responsiveness.

#### Rescue quantification

To express drug-mediated reversal in GSE277437 on a common scale, we computed a Rescue Index (RI) from median Euclidean distances to the frozen Fibrosis centroid: RI = (D_TGF-β1+THZ1_ -D_TGF-β1_)/(D_Untreated_ -D_TGF-β1_). An index of zero indicates no reversal of the TGF-β1-induced ECM shift and 1 indicates a full return to the untreated position.

### 6.8 ECM AGING SCORE

For each of the 180 ADSIG panel ratios, we computed the continuous log-ratio r_ij = log₂(FPKM_{A,i}+1) − log₂(FPKM_{B,i}+1) across all healthy GSE113957 dermal fibroblasts (1–94 y, HGPS excluded) and fitted r_ij ∼ Age_i by ordinary least squares (OLS). Ratios passing Benjamini–Hochberg FDR < 0.05 were retained, and their slopes β_j were cached as age-trend weights. As a sensitivity check, repeating the analysis from read counts for the same 143 samples (recount3, SRP144355) left the phenotype ordering of the aging and HGPS trajectories unchanged. On GSE113957 itself, we computed S^cont and S^sign using the same β-weights as an internal encoding-equivalence check (r = 0.963), confirming that the sign-based encoding retains the ranking structure of the continuous fit on the training data. Because both scores share fitting, selection, and evaluation samples, this correlation validates the encoding, not the aging score’s generalization; the latter is assessed independently on GSE165176 (Section 3.7), which was held out from all fitting steps. S^sign applies these continuous-fit slopes to the discretized sign feature as a convenience weighting scheme, prioritizing consistency with the deployed ECMExplainer sign-encoding architecture overweight-scale purity; we report this as an approximation rather than an equivalence.

The frozen β-weights and ratio panel were then applied, without re-fitting, to GSE165176 (Sendai-OSKM reprogramming of human fibroblasts, 124 samples across 6 donors and multiple timepoints, spanning eight population states: Baseline, Control, ControlIntermed, Failing, FailingIntermed, ReprogrammingIntermed, Reprogramming, and iPSC — with the Reprogramming trajectory enriched by SSEA4⁺ sorting and the Failing/Control trajectory by CD13⁺ sorting). GSE165176 is deposited as log₂-RPM (reads-per-million), not RPKM/FPKM/TPM; RPM is not length-normalized, so unlike RPKM-type units it does not introduce a per-gene length factor that could break sign(log₂-RPM_A − log₂-RPM_B) = sign(true log-CPM difference). For each GSE165176 sample, the same ratios were reconstructed from the author-normalized log₂-RPM matrix and the sign-based score S^sign_i was computed using the GSE113957-derived β-weights. Because β_j is the age-trend slope, S^sign increases with age by construction; a lower score is therefore oriented as younger, justifying the one-sided alternative = ’less’ when testing for rejuvenation. Per-population distributions were visualized against the GSE113957 age-bin reference bands for biological context only, not as a statistical comparator. One-sided Wilcoxon rank-sum tests compared each downstream population to the GSE165176 Baseline (Day 0) distribution within-study; Control and iPSC had fewer than 3 samples with complete age-ratio coverage at the tested endpoints and were excluded from formal testing, along with ControlIntermed, FailingIntermed, and ReprogrammingIntermed, which were shown in trajectory plots for descriptive context only. Reported p-values for the two tested populations (Reprogramming, Failing) were Benjamini–Hochberg adjusted for multiple comparisons. Because the Reprogramming trajectory was SSEA4⁺-sorted and the Failing/Control trajectory CD13⁺-sorted, observed differences in ECM Aging Score partly reflect sort-defined subpopulation composition alongside reprogramming state, and cannot be fully attributed to reprogramming outcome alone.

### 6.9 Model benchmarking

Prediction from all five models [ADSIG-180, POOL-180, ALLGENE-180, All Genes (limma), ECM Genes (limma)] were scored for comparisons. Each gene in the final ADSIG-180, POOL-180 and ALLGENE-180 ratio panels was annotated with its class from the curated ECM genes list (Assembly, Destruction or Signaling), and each ratio was assigned to one of six unordered pair classes (A-D, SIG-SIG, A-A, D-D, A-SIG, D-SIG). All other genes were labeled non-ECM.

#### Null model for pair-class composition

To account for the uneven sizes of the three functional classes, we enumerated all C(711,2) = 252,405 unordered pairs of the POOL ECM universe and counted how many fall into each pair class. This gives the exact expected proportion under random pairing within the same universe.

#### ECM content of the genome-wide panel

ECM representation in ALLGENE-180 was tested by hypergeometric test of the 59 curated ECM genes among the 360 panel genes against the ECM fraction of the 2,000-gene top-variable genes. Pair composition (ECM-ECM, ECM-other, other-other) was tested against the exact random-pairing null for that universe by binomial test with Benjamini-Hochberg correction.

### 6.10 Software

Transcript-level quantification was performed on a Linux server (Ubuntu 24.04 LTS) running R 4.3.3 (Bioconductor 3.18), using Salmon v1.10.2 and tximport v1.30.0. All downstream statistical analyses, model training, and figure generation were performed on a local workstation running R 4.3.3 (Bioconductor 3.18). Key packages: Salmon v1.10.2, tximport v1.30, MASS v7.3, limma v3.58, glmnet v4.1 (sensitivity analyses only), ggplot2 v3.5, patchwork v1.3, ComplexHeatmap v2.20, dplyr/tidyr (tidyverse 2.0), pROC v1.18, reshape2 v1.4.4, GEOquery v2.70, AnnotationDbi v1.64, org.Hs.eg.db v3.18, scatterplot3d v0.3, ggrepel v0.9, scales v1.3, and viridis v0.6.

### 6.11 Use of AI tools

Large language model assistants (Claude, ChatGPT, Perplexity) were used to help write and debug the R, Python and shell scripts for data processing, model training, validation and figure generation, to assemble and cross-check some supplementary tables from author-generated result files and to proofread the manuscript. All AI-generated code, tables and text edits were checked and revised by the authors, who designed the study, interpreted the results and take full responsibility for the content. No AI tool was used to generate data or images.

## 7 Data and code availability

ECMExplainer is available at https://github.com/avelar-ageing/ecmexplainer under the MIT licence; the version used in this study is release v1.0.0. The repository contains the R implementation of ratio encoding, panel selection, training and projection, the three trained panels (ADSIG-180, POOL-180 and ALLGENE-180), the curated matrisome gene list, Supplementary Tables 1–6, and a Shiny application that classifies

user-supplied expression data, retrains panels on user-defined labels and gene lists, and computes the ECM Aging Score. Included tests reproduce every validation prediction in Supplementary Table 6 (R 4.4, org.Hs.eg.db 3.18). The application runs locally in R or Docker, and a public instance is available at https://apps.avelar.science/ecmexplainer/.

## 9 Author Contributions

E.S.: Conceptualization, Methodology, Formal analysis, Data curation, Investigation, Visualization, Writing – original draft, review and editing. R.A.A: Conceptualization, Methodology, Writing – review and editing. H.F: Supervision, Resources, Funding acquisition, Writing – review and editing.

## 11 Acknowledgements

We thank members of the Fujiwara laboratory for their helpful discussion throughout the project, and the authors of the publicly available datasets analyzed here for making their data accessible.

## Supplementary Figures

**Supplementary Figure 1:**
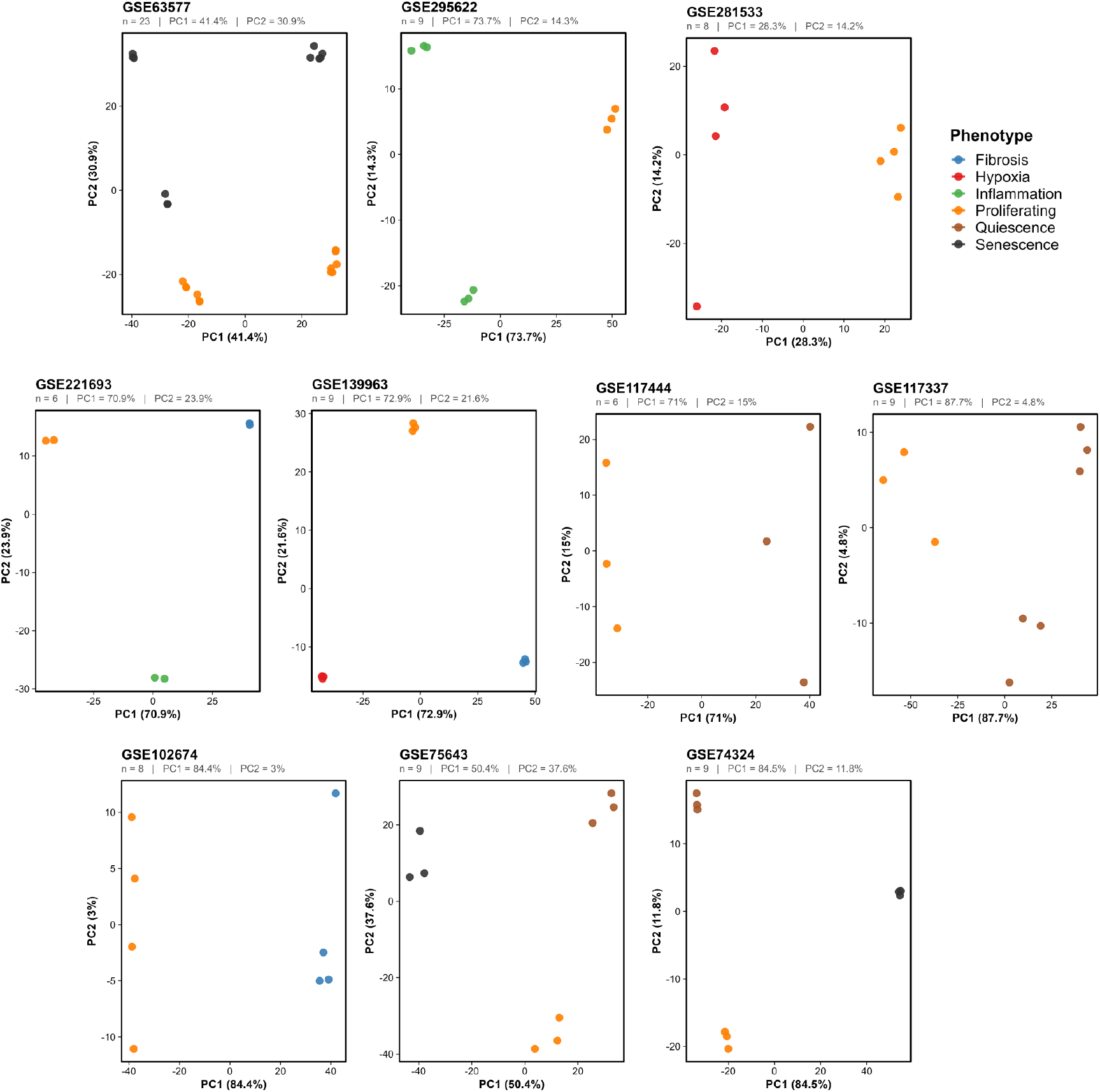
PCA of the 96 sample training samples. All show clear phenotype separation.

**Supplementary Figure 2:**
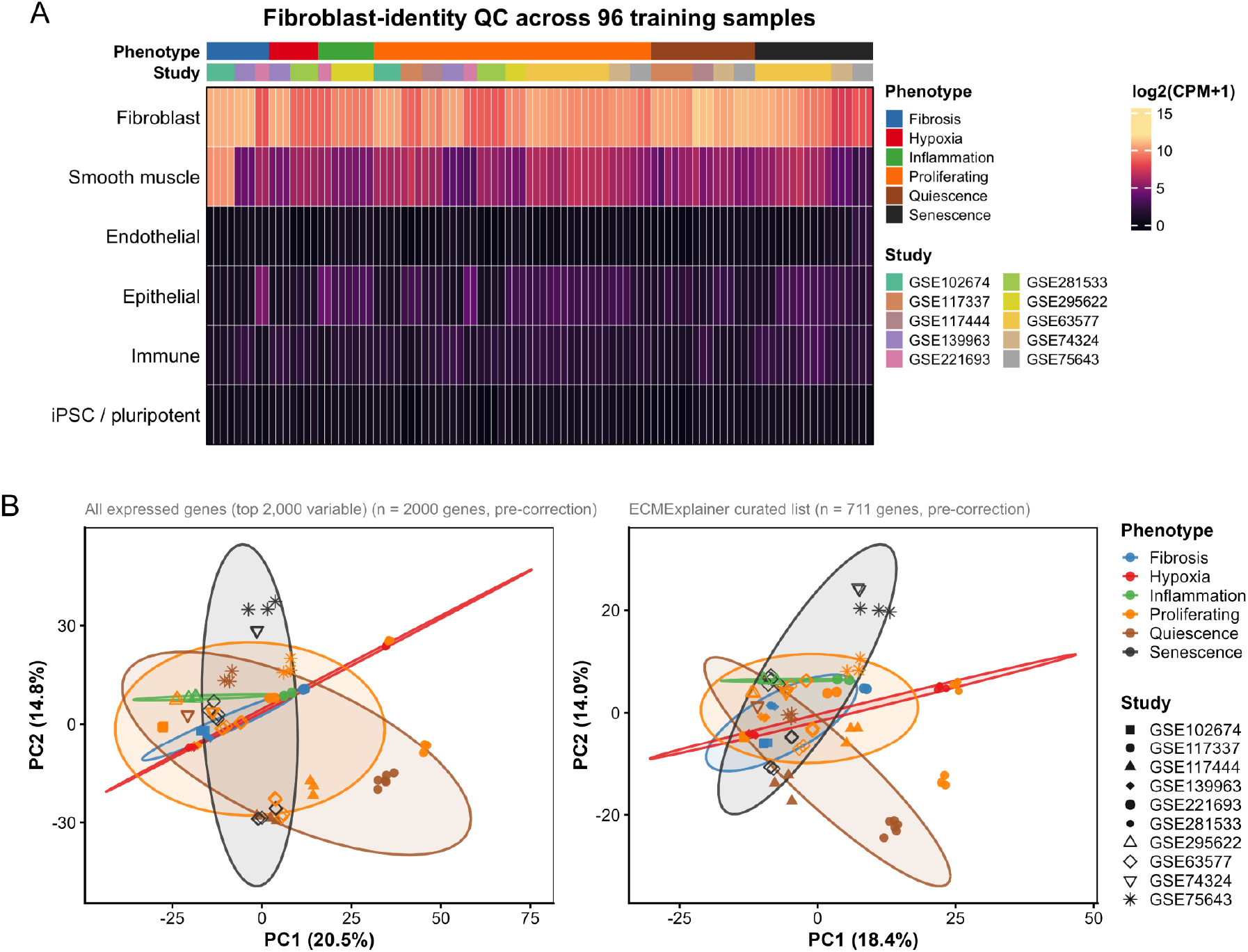
(A) Fibroblast-identity QC across the 96 training samples. Mean log2(CPM+1) expression of canonical fibroblast markers was compared against epithelial, endothelial, immune, smooth-muscle and pluripotent-stem-cell lineage panels for each sample. Colored bars above the heatmap indicate phenotype (top) and source study (bottom); samples are ordered by phenotype, then study, then fibroblast-panel score. (B) Global PCA of the 96 training samples before batch correction for (Left) 2000 variable genes and (Right) ECMExplainer curated gene list genes, colored by phenotype and shaped by source study. Strong study-driven clustering is evident on PC1 and PC2, consistent with technical variation across sequencing pipelines and culture conditions.

**Supplementary Figure 3:**
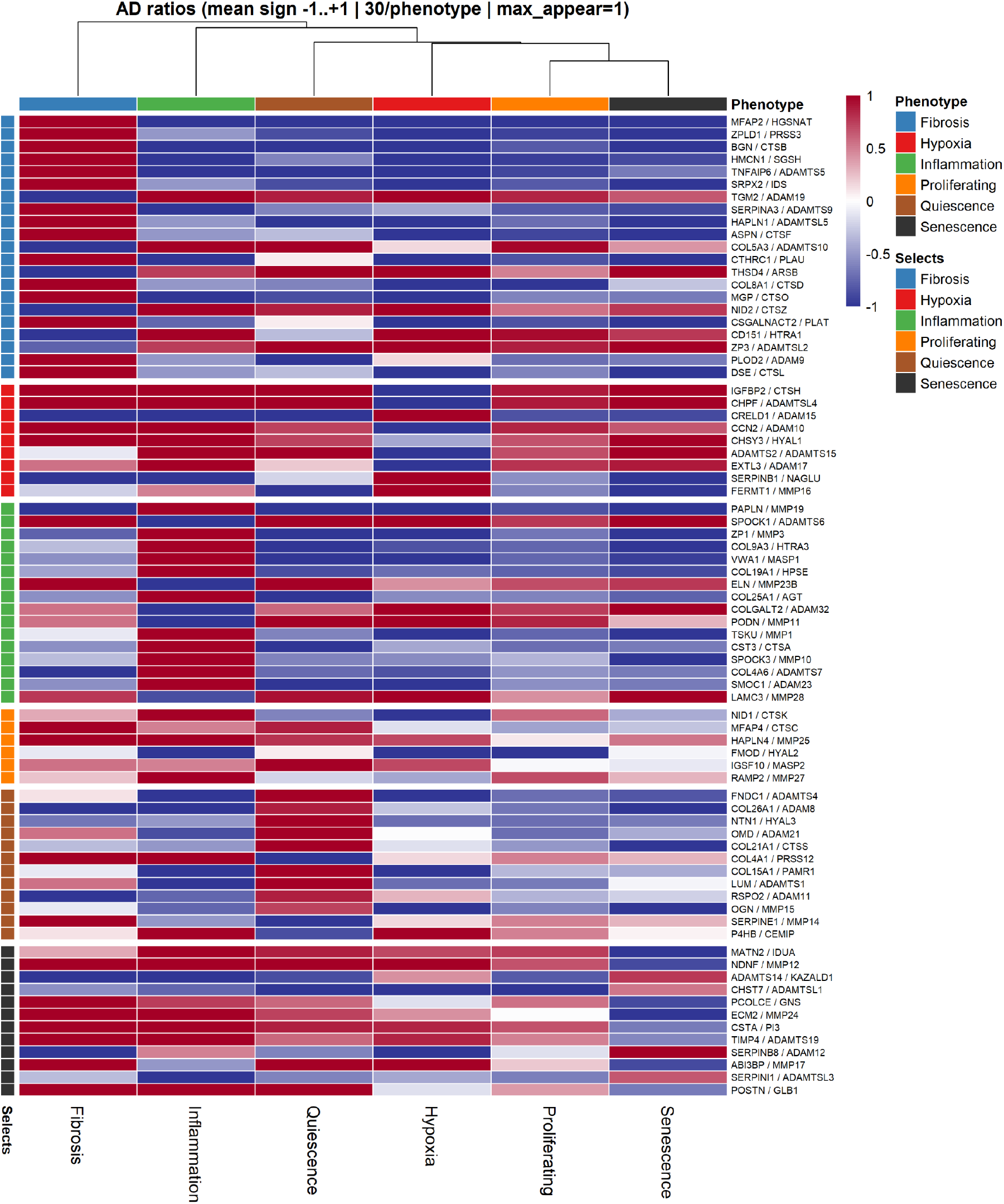
Phenotype clustering of Assembly/Destruction (A/D) ratios. Heatmap of the mean sign (−1 to +1) of the 76 A/D ratios comprising the ECMExplainer panel, averaged across all samples within each phenotype, with both phenotypes (columns) and ratios (rows) hierarchically clustered (Euclidean distance, complete linkage). Row annotations ("Selects") indicate the phenotype for which each ratio was originally selected during the greedy quota procedure; row blocks are grouped by selecting phenotype. Color key denotes mean ratio sign: dark red = consistently gene A > gene B (+1) across the phenotype’s samples, dark blue = consistently gene A < gene B (−1), white = mixed/no consistent sign (0).

**Supplementary Figure 4:**
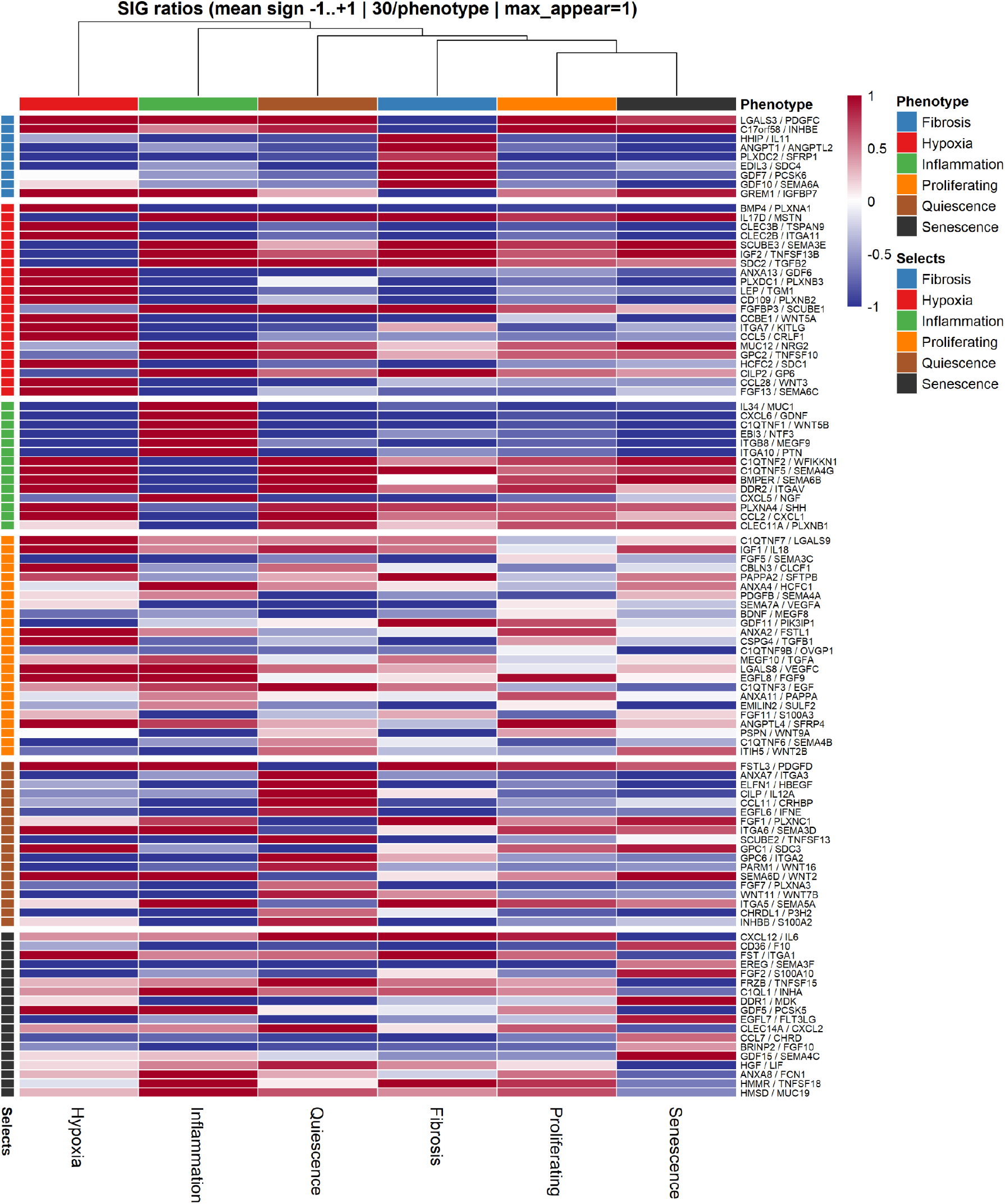
Phenotype-level clustering of Signaling/Signaling (SIG) ratios. Heatmap of the mean sign (−1 to +1) of the 104 SIG ratios comprising the ECMExplainer panel, averaged across all samples within each phenotype, with both phenotypes (columns) and ratios (rows) hierarchically clustered (Euclidean distance, complete linkage). Row annotations ("Selects") indicate the phenotype for which each ratio was originally selected during the greedy quota procedure; row blocks are grouped by selecting phenotype. Color key denotes mean ratio sign: dark red = consistently gene A > gene B (+1) across the phenotype’s samples, dark blue = consistently gene A < gene B (−1), white = mixed/no consistent sign (0).

**Supplementary Figure 5:**
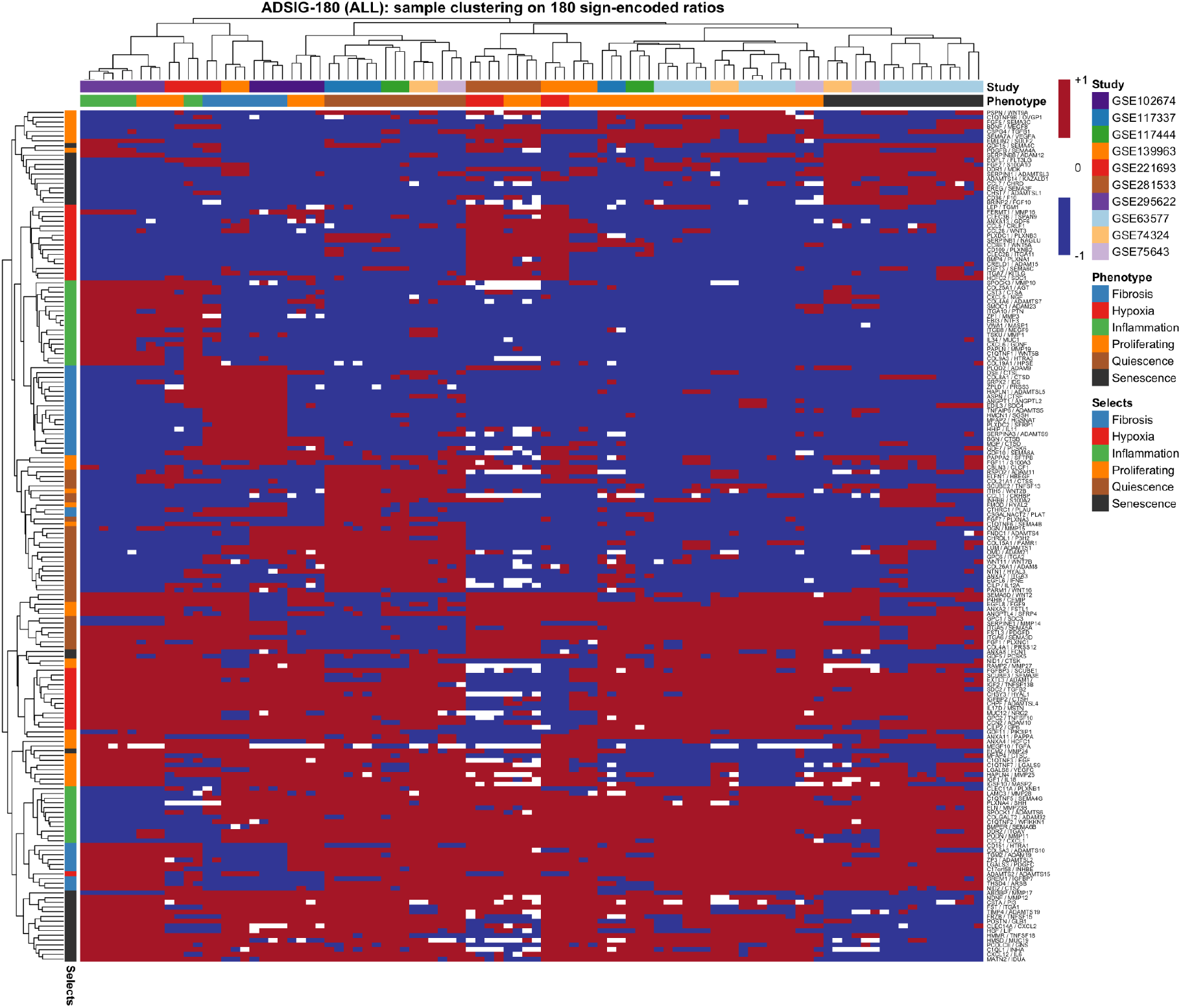
Sample-level hierarchical clustering on the ADSIG-180 sign-encoded ratio matrix. Heatmap of the sign (−1, 0, +1) of all 180 gene-pair log-ratios comprising the final ADSIG classifier panel, with both samples (columns) and ratios (rows) hierarchically clustered (Euclidean distance, complete linkage). Column annotations indicate each sample’s source study and assigned phenotype; row annotations ("Selects") indicate the phenotype for which each ratio was originally selected during the greedy quota procedure. Color key denotes ratio sign: dark red = +1 (gene A > gene B), dark blue = −1 (gene A < gene B), white = 0 (tie).

**Supplementary Figure 6:**
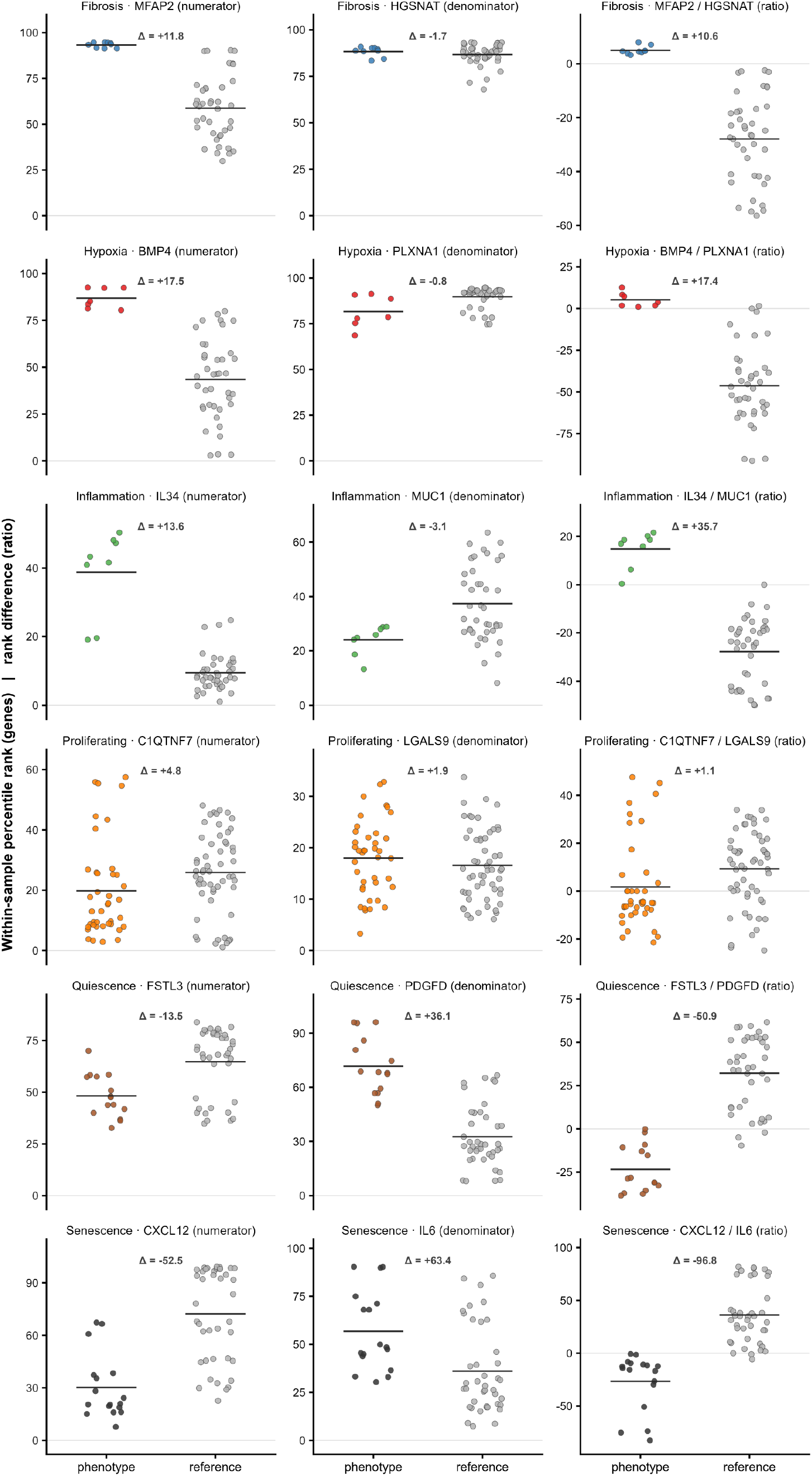
Expression behavior of the leading discriminative ratio for each phenotype. For each leading discriminative ratio, each point shows one sample’s value for the numerator gene, the denominator gene, and the difference between them, which is the quantity the classifier reads. Colored points represent the selecting phenotype. Grey points are the within-study reference: Proliferating samples for the five non-proliferating phenotypes, and all non-proliferating samples for the Proliferating panel. Gene values are within-sample percentile ranks, computed for each sample across (see Methods). Δ gives the pooled shift (phenotype minus reference, in percentile points). Equivalent shifts with 95% confidence intervals, behavior class and reliability for all 180 ratios are provided in Supplementary Table 3.

**Supplementary Figure 7:**
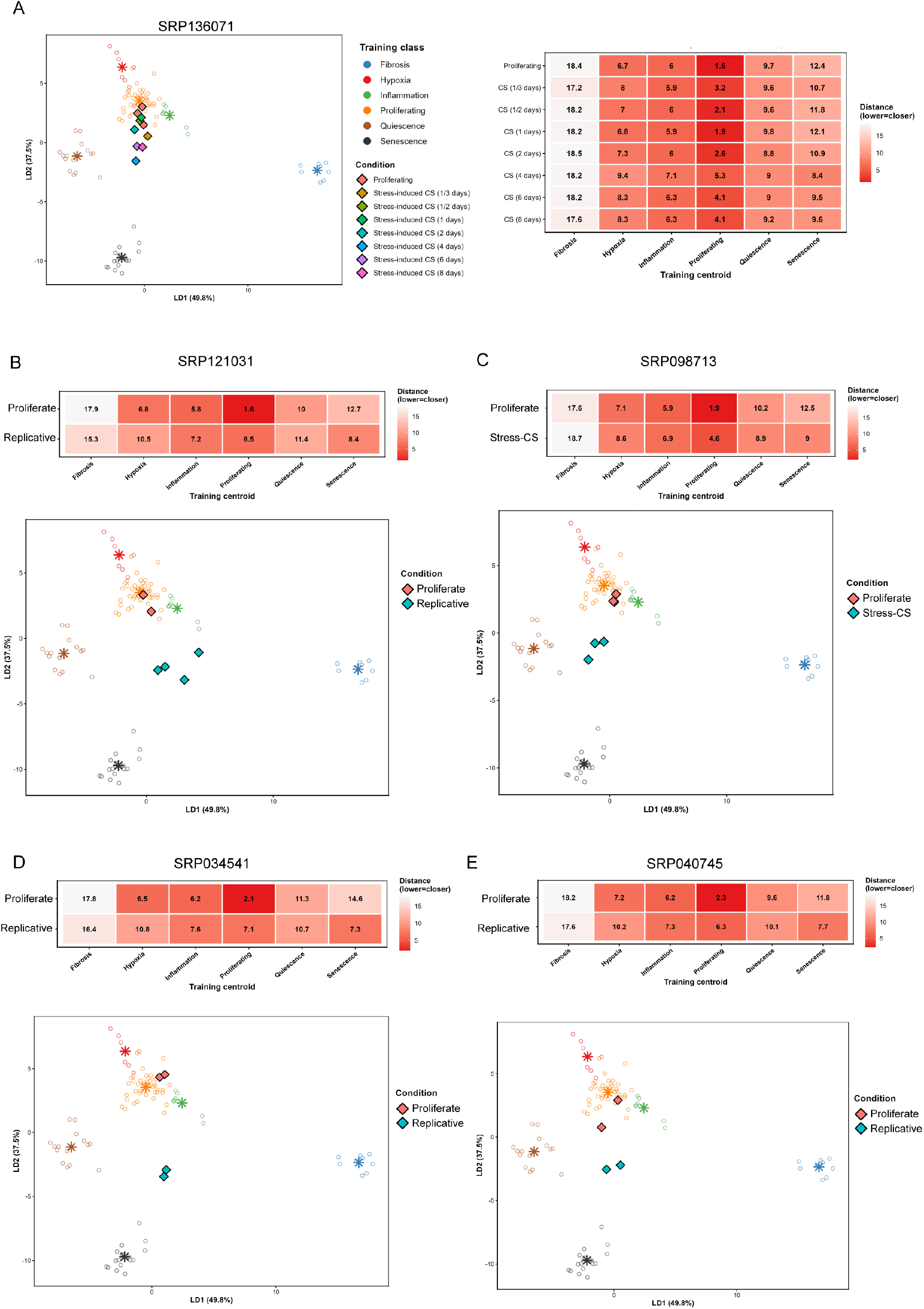
Additional Senescence validation: Five independent datasets: (A) SRP136071, (B) SRP121031, (C) SRP098713, (D) SRP034541, (E) SRP040745 projected onto the ECMExplainer LDA space.

**Supplementary Figure 8:**
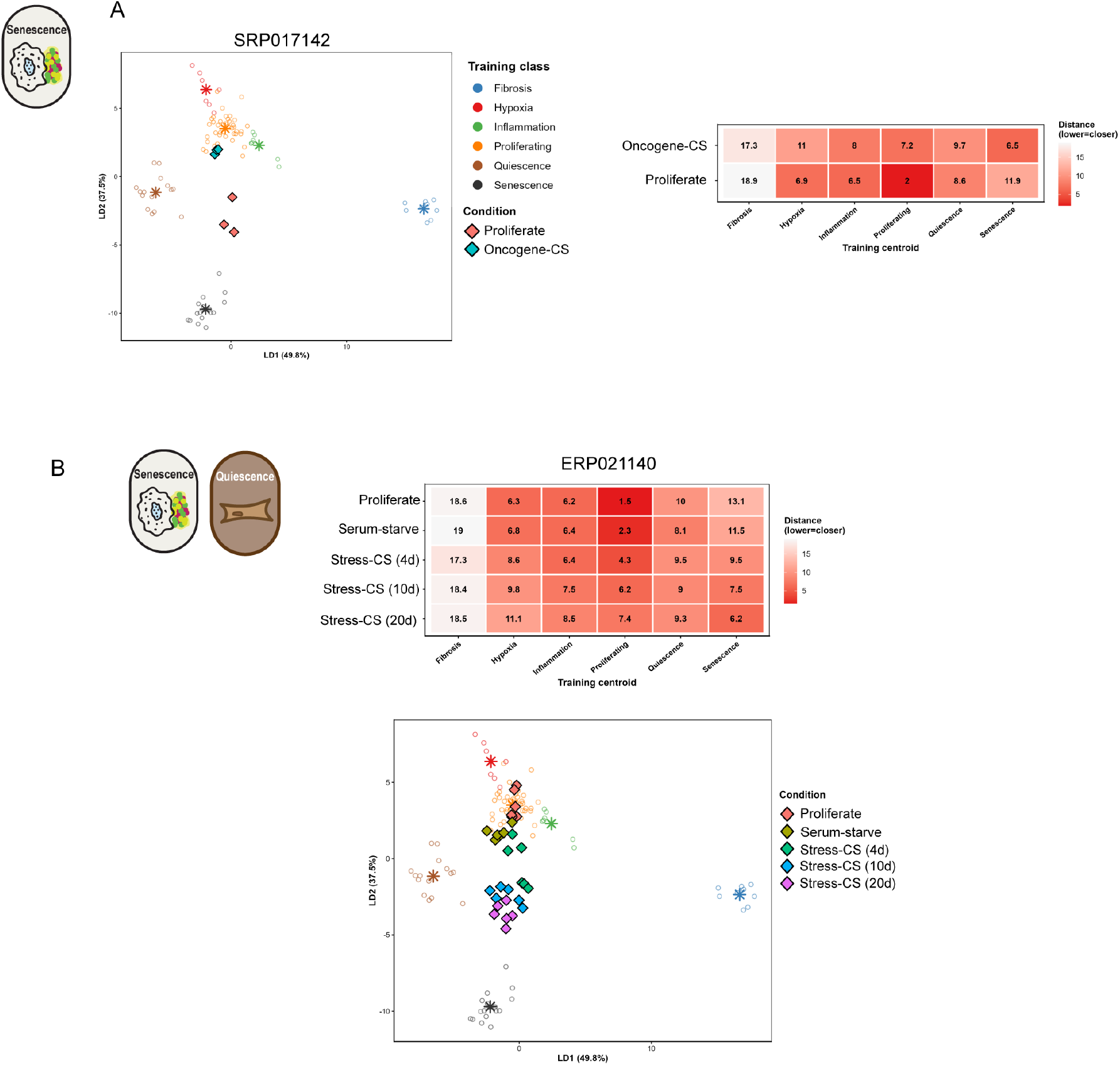
Additional oncogene-induced senescence and combined cellular senescence and quiescence validation. Two datasets: (A) SRP017142, (B) ERP021140 projected onto the ECMExplainer LDA space. ERP021140 includes paired quiescence and senescence arms, demonstrating that the ECMExplainer correctly resolves the two non-proliferative states in the same experimental system.

**Supplementary Figure 9:**
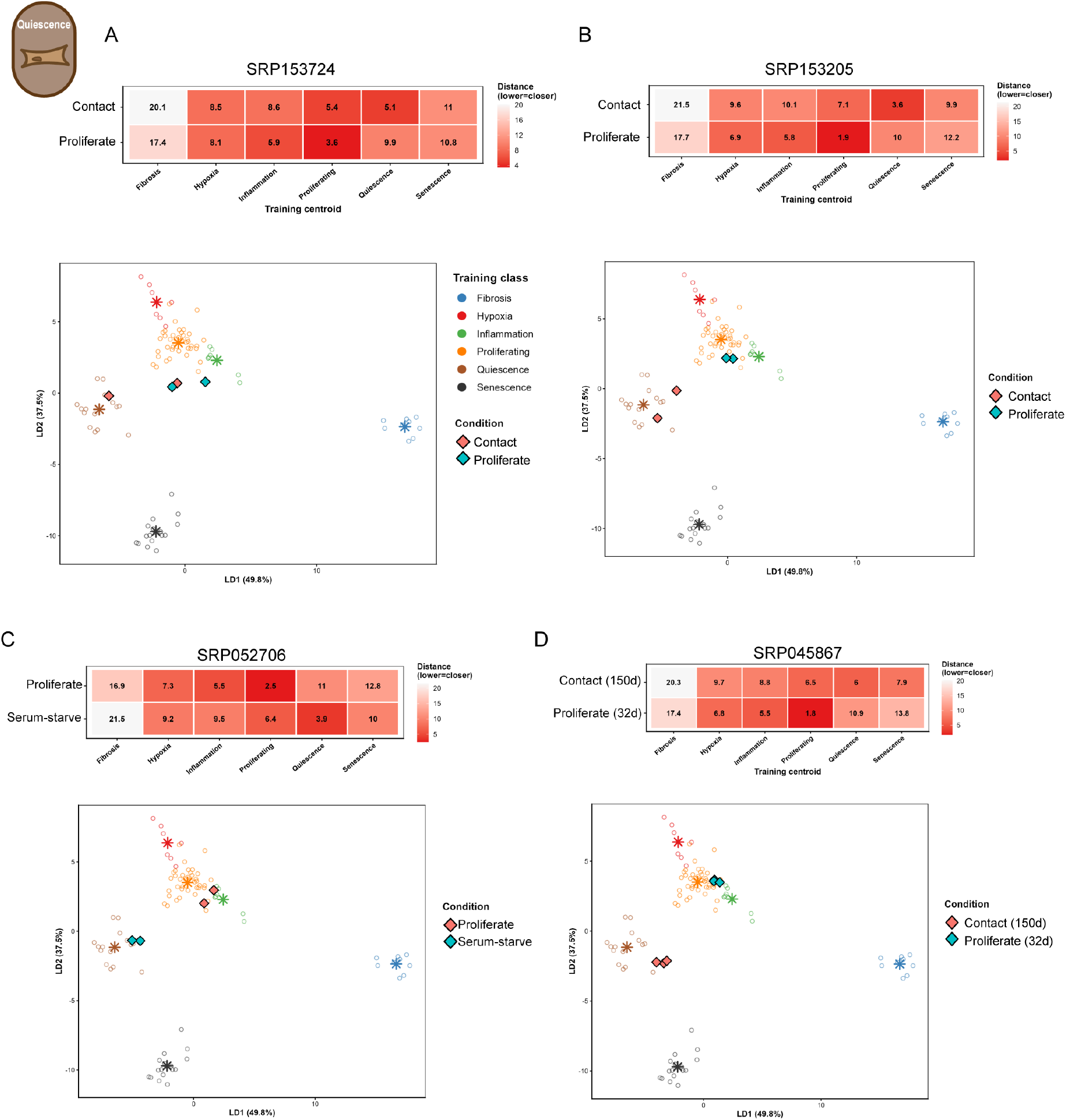
Additional quiescence validation. Four independent contact-inhibited or serum-starved cellular quiescence (CQ) datasets: (A) SRP153724, (B) SRP153205, (C) SRP052706, (D) SRP045867 projected onto the ECMExplainer LDA space; CQ samples localize to the Quiescence centroid and proliferating controls to the Proliferating centroid. Median Euclidean distance heatmaps shown below each LDA panel.

**Supplementary Figure 10:**
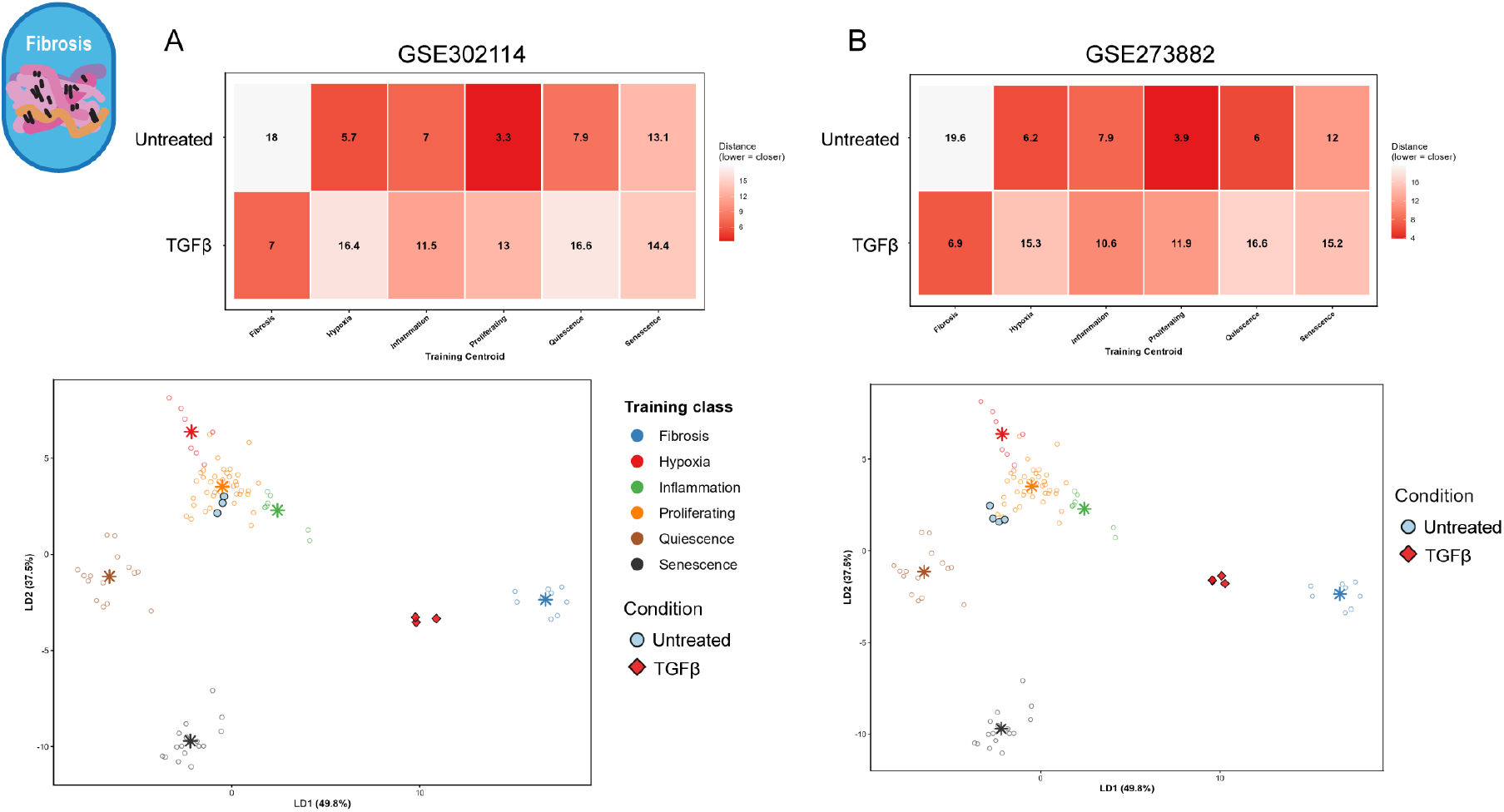
Fibrosis validation on TGFβ-stimulated fibroblasts. LDA projections (upper row) and median Euclidean distance heatmaps (lower row) for two independent fibrosis datasets: (A) GSE302114 (untreated vs TGFβ-treated) and (B) GSE273882 (untreated vs TGFβ-treated).

**Supplementary Figure 11:**
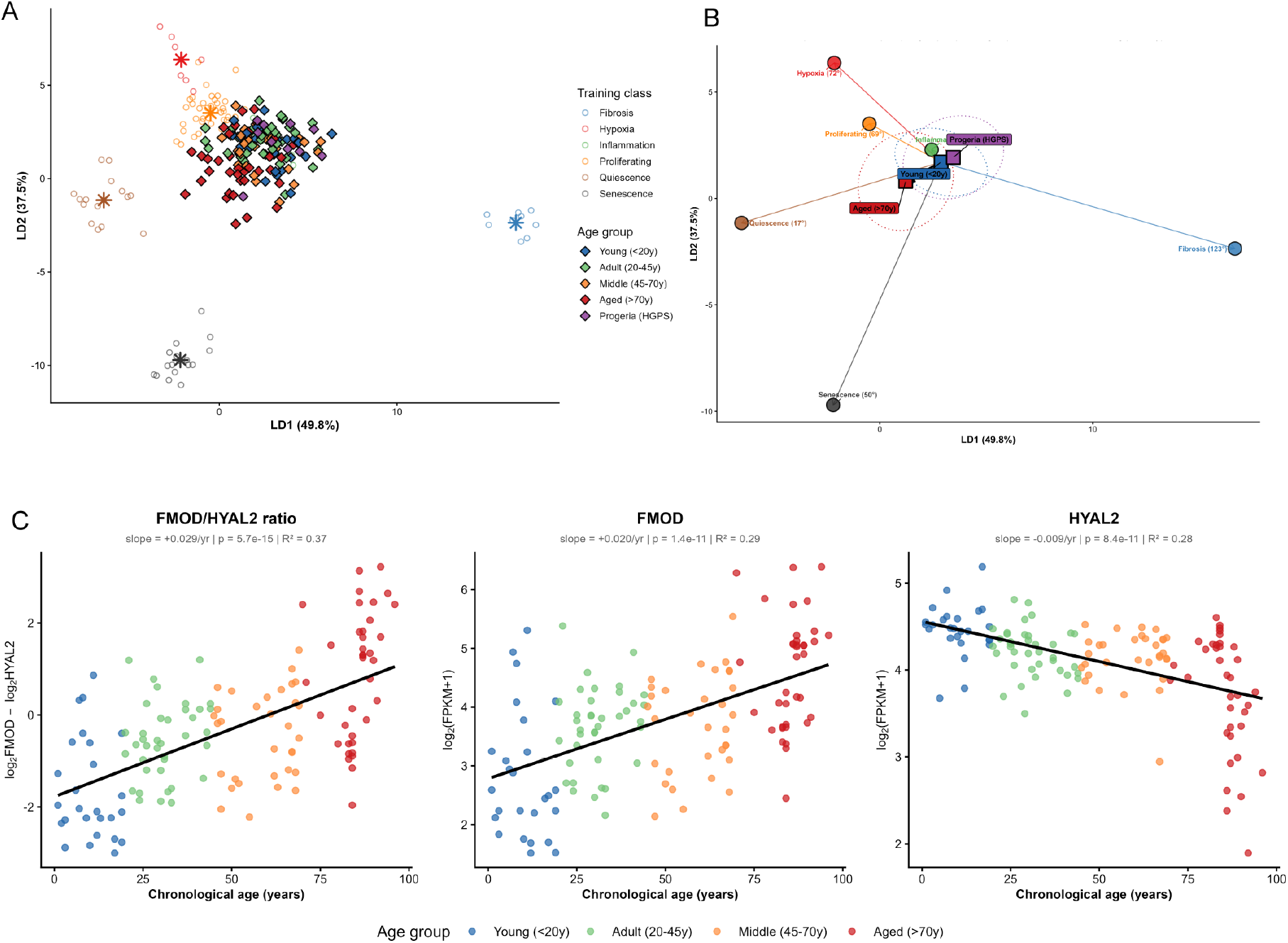
(A) LDA projection for GSE113957. (B) Derivation of protractor vectors. Squares denote Young, Aged, and HGPS group centroids; circles denote training phenotypes. The solid arrow defines the Young-to-Aged 0° reference trajectory and the dashed arrow the Young-to-HGPS trajectory. Angles shown are calculated in the full five-dimensional space, while LD1-LD2 are displayed for visualization. (C) The directional split of those significant ratios into up-versus down-regulated with age. Of 180 ratios, 96 (53%) change significantly with donor age (linear regression of each continuous ratio on age, Benjamini–Hochberg FDR < 0.05).

## Supplementary Table information

Supplementary Table 1: ECMExplainer curated gene list and functional module assignment

Supplementary Table 2: Training cohort, study support and lineage-quality control

Supplementary Table 3: ADSIG-180 ratio panel and constituent gene behavior

Supplementary Table 4: Age-associated ratios and ECM Aging Score

Supplementary Table 5: POOL-180 and ALLGENE-180 sweep and ratio lists

Supplementary Table 6: Benchmark-panel composition and comparisons

## References

1. Naba, A. et al. The extracellular matrix: Tools and insights for the ‘omics’ era. Matrix Biol 49, 10–24 (2016).

2. Naba, A., et al. The Matrisome: In Silico Definition and In Vivo Characterization by Proteomics of Normal and Tumor Extracellular Matrices*. Molecular & Cellular Proteomics 11, M111.014647 (2012).

3. Shao, X. et al. MatrisomeDB 2.0: 2023 updates to the ECM-protein knowledge database. Nucleic Acids Res 51, D1519–D1530 (2023).

4. Omachi, K. & Fujiwara, H. Exploring basement membrane dynamics through cross-scale imaging, manipulation, and molecular mapping. Matrix Biol 141, 67– 81 (2025).

5. Kroemer, G. et al. From geroscience to precision geromedicine: Understanding and managing aging. Cell 188, 2043–2062 (2025).

6. Park, J. Y. C. et al. Strategic outline of interventions targeting extracellular matrix for promoting healthy longevity. Am J Physiol Cell Physiol 325, C90–C128 (2023).

7. Coenen, L., Molière, A., Lehallier, B., Middeldorp, J. & Ewald, C. Y. An Extracellular Matrix Aging Clock Based on Circulating Matrisome Proteins Predicts Biological Aging and Disease. Aging Cell 25, e70474 (2026).

8. Coppé, J.-P., Desprez, P.-Y., Krtolica, A. & Campisi, J. The senescence-associated secretory phenotype: the dark side of tumor suppression. Annu Rev Pathol 5, 99–118 (2010).

9. Selman, M. & Pardo, A. Fibroageing: An ageing pathological feature driven by dysregulated extracellular matrix-cell mechanobiology. Ageing Research Reviews 70, 101393 (2021).

10. Hernández-Bellido, N., Perramon-Güell, A. & Anerillas, C. The aging extracellular matrix as a missing link in senescent cell accumulation and persistence. Nat Aging https://doi.org/10.1038/s43587-026-01127-w (2026) doi:10.1038/s43587-026-01127-w.

11. Kumar, R. K. Morphological methods for assessment of fibrosis. Methods Mol Med 117, 179–188 (2005).

12. Zhang, Q., An, Z.-Y., Jiang, W., Jin, W.-L. & He, X.-Y. Collagen code in tumor microenvironment: Functions, molecular mechanisms, and therapeutic implications. Biomed Pharmacother 166, 115390 (2023).

13. Nag, R. et al. Collagen disorder architecture features are associated with clinical, molecular, genetic factors and survival outcomes in colon cancer. NPJ Precis Oncol 9, 304 (2025).

14. Statzer, C. et al. Youthful and age-related matreotypes predict drugs promoting longevity. Aging Cell 20, e13441 (2021).

15. Petrov, P. B., Considine, J. M., Izzi, V. & Naba, A. Matrisome AnalyzeR -a suite of tools to annotate and quantify ECM molecules in big datasets across organisms. J Cell Sci 136, jcs261255 (2023).

16. Lamba, R., Paguntalan, A. M., Petrov, P. B., Naba, A. & Izzi, V. MatriCom, a single-cell RNA-sequencing data mining tool to infer cell-extracellular matrix interactions. J Cell Sci 138, jcs263927 (2025).

17. Oshinjo, A., Chen, D., Petrov, P., Izzi, V. & Naba, A. MatriSpace: Identification and visualization of spatially resolved ECM gene expression patterns in health and disease. 2026.04.26.720198 Preprint at 10.64898/2026.04.26.720198 (2026).

18. Liberzon, A. et al. The Molecular Signatures Database (MSigDB) hallmark gene set collection. Cell Syst 1, 417–425 (2015).

19. Calcinotto, A. et al. Cellular Senescence: Aging, Cancer, and Injury. Physiological Reviews 99, 1047–1078 (2019).

20. Li, X. et al. Inflammation and aging: signaling pathways and intervention therapies. Signal Transduct Target Ther 8, 239 (2023).

21. Buechler, M. B. et al. Cross-tissue organization of the fibroblast lineage. Nature 593, 575–579 (2021).

22. Muhl, L. et al. Single-cell analysis uncovers fibroblast heterogeneity and criteria for fibroblast and mural cell identification and discrimination. Nat Commun 11, 3953 (2020).

23. Tabula Sapiens Consortium* et al. The Tabula Sapiens: A multiple-organ, single-cell transcriptomic atlas of humans. Science 376, eabl4896 (2022).

24. Owens, G. K., Kumar, M. S. & Wamhoff, B. R. Molecular regulation of vascular smooth muscle cell differentiation in development and disease. Physiol Rev 84, 767–801 (2004).

25. Takahashi, K. et al. Induction of pluripotent stem cells from adult human fibroblasts by defined factors. Cell 131, 861–872 (2007).

26. Hinz, B. et al. Recent developments in myofibroblast biology: paradigms for connective tissue remodeling. Am J Pathol 180, 1340–1355 (2012).

27. Yao, G. Modelling mammalian cellular quiescence. Interface Focus 4, 20130074 (2014).

28. Geman, D., d’Avignon, C., Naiman, D. Q. & Winslow, R. L. Classifying gene expression profiles from pairwise mRNA comparisons. Stat Appl Genet Mol Biol 3, Article19 (2004).

29. Tan, A. C., Naiman, D. Q., Xu, L., Winslow, R. L. & Geman, D. Simple decision rules for classifying human cancers from gene expression profiles. Bioinformatics 21, 3896–3904 (2005).

30. Koenitzer, J., Wu, H., Brody, S. L., Mecham, R. P. & Humphreys, B. D. Microfibril-Associated Glycoproteins Attenuate Pulmonary Fibrosis in Mice. Am J Respir Crit Care Med 203, A4427 (2021).

31. Zhang, W. et al. Hepatic stellate cell-derived microfibrillar-associated protein 2 prevents liver fibrosis by regulating extracellular matrix and inflammation. Theranostics 15, 4033–4053 (2025).

32. Frank, D. B. et al. Bone morphogenetic protein 4 promotes pulmonary vascular remodeling in hypoxic pulmonary hypertension. Circ Res 97, 496–504 (2005).

33. Cao, H. et al. PDGFD: A Dual-Function Regulator That Maintains Myoblast Pool and Fuels Myogenic Differentiation. Current Issues in Molecular Biology 48, 322 (2026).

34. Baghdadi, M. et al. Interleukin-34, a comprehensive review. Journal of Leukocyte Biology 104, 931–951 (2018).

35. Dhar, P. & McAuley, J. The Role of the Cell Surface Mucin MUC1 as a Barrier to Infection and Regulator of Inflammation. Front. Cell. Infect. Microbiol. 9, (2019).

36. Acosta, J. C. et al. A complex secretory program orchestrated by the inflammasome controls paracrine senescence. Nat Cell Biol 15, 978–990 (2013).

37. Coppé, J.-P. et al. Senescence-associated secretory phenotypes reveal cell-nonautonomous functions of oncogenic RAS and the p53 tumor suppressor. PLoS Biol 6, 2853–2868 (2008).

38. Georgilis, A. et al. PTBP1-Mediated Alternative Splicing Regulates the Inflammatory Secretome and the Pro-tumorigenic Effects of Senescent Cells. Cancer Cell 34, 85–102.e9 (2018).

39. Aarts, M. et al. Coupling shRNA screens with single-cell RNA-seq identifies a dual role for mTOR in reprogramming-induced senescence. Genes Dev 31, 2085–2098 (2017).

40. Wang, R. et al. Rapamycin inhibits the secretory phenotype of senescent cells by a Nrf2-independent mechanism. Aging Cell 16, 564–574 (2017).

41. Azar, A., Lawrence, I., Jofre, S., Mell, J. & Sell, C. Distinct patterns of gene expression in human cardiac fibroblasts exposed to rapamycin treatment or methionine restriction. Ann N Y Acad Sci 1418, 95–105 (2018).

42. Schafer, S. et al. IL-11 is a crucial determinant of cardiovascular fibrosis. Nature 552, 110–115 (2017).

43. Ma, H.-Y. et al. Targeting CDK7/12/13 functional synergism reverses myofibroblast activation and ameliorates lung fibrosis. iScience 28, 112778 (2025).

44. Sen, P. et al. Histone Acetyltransferase p300 Induces De Novo Super-Enhancers to Drive Cellular Senescence. Mol Cell 73, 684–698.e8 (2019).

45. Rai, T. S. et al. HIRA orchestrates a dynamic chromatin landscape in senescence and is required for suppression of neoplasia. Genes Dev 28, 2712– 2725 (2014).

46. Alspach, E. et al. p38MAPK plays a crucial role in stromal-mediated tumorigenesis. Cancer Discov 4, 716–729 (2014).

47. Baar, M. P. et al. Targeted Apoptosis of Senescent Cells Restores Tissue Homeostasis in Response to Chemotoxicity and Aging. Cell 169, 132–147.e16 (2017).

48. Zhao, Z. et al. Dynamic transcriptome profiling in DNA damage-induced cellular senescence and transient cell-cycle arrest. Genomics 112, 1309–1317 (2020).

49. Gorgoulis, V. et al. Cellular Senescence: Defining a Path Forward. Cell 179, 813– 827 (2019).

50. Herranz, N. & Gil, J. Mechanisms and functions of cellular senescence. J Clin Invest 128, 1238–1246 (2018).

51. Neyret-Kahn, H. et al. Sumoylation at chromatin governs coordinated repression of a transcriptional program essential for cell growth and proliferation. Genome Res 23, 1563–1579 (2013).

52. Hernandez-Segura, A. et al. Unmasking Transcriptional Heterogeneity in Senescent Cells. Curr Biol 27, 2652–2660.e4 (2017).

53. Mitra, M., Ho, L. D. & Coller, H. A. An In Vitro Model of Cellular Quiescence in Primary Human Dermal Fibroblasts. Methods Mol Biol 1686, 27–47 (2018).

54. Mitra, M. et al. Alternative polyadenylation factors link cell cycle to migration. Genome Biol 19, 176 (2018).

55. Trost, B. et al. Concordance between RNA-sequencing data and DNA microarray data in transcriptome analysis of proliferative and quiescent fibroblasts. R Soc Open Sci 2, 150402 (2015).

56. Marthandan, S., Priebe, S., Hemmerich, P., Klement, K. & Diekmann, S. Long-Term Quiescent Fibroblast Cells Transit into Senescence. PLOS ONE 9, e115597 (2014).

57. Meliton, A. Y. et al. Mitochondrial one-carbon metabolism is required for TGF-β-induced glycine synthesis and fibrotic responses. Nat Commun 16, 9250 (2025).

58. Shin, K. W. D. et al. mTOR signaling regulates multiple metabolic pathways in human lung fibroblasts after TGF-β and in pulmonary fibrosis. Am J Physiol Lung Cell Mol Physiol 328, L215–L228 (2025).

59. Fleischer, J. G. et al. Predicting age from the transcriptome of human dermal fibroblasts. Genome Biol 19, 221 (2018).

60. de Magalhães, J. P., Curado, J. & Church, G. M. Meta-analysis of age-related gene expression profiles identifies common signatures of aging. Bioinformatics 25, 875–881 (2009).

61. Saul, D. et al. A new gene set identifies senescent cells and predicts senescence-associated pathways across tissues. Nat Commun 13, 4827 (2022).

62. Biran, A. et al. Quantitative identification of senescent cells in aging and disease. Aging Cell 16, 661–671 (2017).

63. Evans, C. H. On the aging of organisms and their cells. Med Hypotheses 5, 53– 66 (1979).

64. Ocampo, A. et al. In Vivo Amelioration of Age-Associated Hallmarks by Partial Reprogramming. Cell 167, 1719–1733.e12 (2016).

65. Simpson, D. J., Olova, N. N. & Chandra, T. Cellular reprogramming and epigenetic rejuvenation. Clin Epigenet 13, 170 (2021).

66. Lu, J. Y. et al. Prevalent mesenchymal drift in aging and disease is reversed by partial reprogramming. Cell 188, 5895–5911.e17 (2025).

67. Gill, D. et al. Multi-omic rejuvenation of human cells by maturation phase transient reprogramming. Elife 11, e71624 (2022).

68. Kirschner, K., Rattanavirotkul, N., Quince, M. F. & Chandra, T. Functional heterogeneity in senescence. Biochem Soc Trans 48, 765–773 (2020).

69. Marthandan, S. et al. Conserved Senescence Associated Genes and Pathways in Primary Human Fibroblasts Detected by RNA-Seq. PLoS One 11, e0154531 (2016).

70. Marthandan, S. et al. Similarities in Gene Expression Profiles during In Vitro Aging of Primary Human Embryonic Lung and Foreskin Fibroblasts. Biomed Res Int 2015, 731938 (2015).

71. Tasdemir, N. et al. BRD4 Connects Enhancer Remodeling to Senescence Immune Surveillance. Cancer Discov 6, 612–629 (2016).

72. Lenain, C. et al. Massive reshaping of genome-nuclear lamina interactions during oncogene-induced senescence. Genome Res 27, 1634–1644 (2017).

73. Selvarajah, B. et al. mTORC1 amplifies the ATF4-dependent de novo serine-glycine pathway to supply glycine during TGF-β1-induced collagen biosynthesis. Sci Signal 12, eaav3048 (2019).

74. Lee, H. N. et al. RECK isoforms have opposing effects on cell migration. Mol Biol Cell 29, 1825–1838 (2018).

75. Senavirathna, L. K., Huang, C., Pushparaj, S., Xu, D. & Liu, L. Hypoxia and transforming growth factor β1 regulation of long non-coding RNA transcriptomes in human pulmonary fibroblasts. Physiol Rep 8, e14343 (2020).

76. Alexanian, M. et al. Chromatin Remodeling Drives Immune-Fibroblast Crosstalk in Heart Failure Pathogenesis. bioRxiv 2023.01.06.522937 (2023) doi:10.1101/2023.01.06.522937.

77. Suzuki, Y. et al. Transcriptome Analysis of Fibroblasts in Hypoxia-Induced Vascular Remodeling: Functional Roles of CD26/DPP4. Int J Mol Sci 25, 12599 (2024).

78. Ghosh, N., Gupta, R. K., Jung, J., Fung, K. & Croft, M. LIGHT in combination with IL-13 or IL-17 drives inflammatory transcriptional signatures in human pulmonary fibroblasts relevant for human lung disease. Immunohorizons 9, vlaf042 (2025).

79. Patro, R., Duggal, G., Love, M. I., Irizarry, R. A. & Kingsford, C. Salmon provides fast and bias-aware quantification of transcript expression. Nat Methods 14, 417– 419 (2017).

80. Soneson, C., Love, M. I. & Robinson, M. D. Differential analyses for RNA-seq: transcript-level estimates improve gene-level inferences. F1000Res 4, 1521 (2015).

